# Behavior- and Cell Type-Specific Cortico-Striatal Activity Decoupling in a Parkinson’s Disease-Like Mouse Model

**DOI:** 10.1101/2024.08.13.607859

**Authors:** Xu-Ran Yao, Yang Liu, Wei-Tong Zheng, Mao-Qing Huang, Kai-Wen He

## Abstract

Inter-brain region activity coupling is essential for enabling coordinated neural communication, facilitating complex brain processes including motor behaviors. In Parkinson’s Disease (PD) patients, cortico-striatal decoupling is widely reported while its onset and cellular mechanism remain largely unclear. Using dual-site fiber photometry and Cre transgenic mouse lines, we examined activity coordination between M1 cortex and dorsal striatum (cortico-striatal coupling) with cell-type resolution. This method identifies motor behavior-specific coupling patterns with different contribution from striatal D1R- and D2R-expressing medium spiny neurons (MSNs). In an α-synuclein preformed fibrils (PFF) induced PD-like mouse model, cortico-striatal coupling associated with digging behavior is selectively disrupted as early as two months post-induction, whose progressive deterioration correlates with later-onset behavioral deficits. Optogenetic disruption of cortico-striatal coupling is sufficient to induce digging deficits in wild-type mice. Furthermore, such decoupling is mainly mediated by impaired D1R MSNs, which can be rescued by D1 receptor activation or L-DOPA. These findings demonstrate that early-onset, behavior- and cell type-specific cortico-striatal decoupling emerges early during the development of PD-like symptoms.

## INTRODUCTION

Motor behavior relies on dynamic, coordinated activity across distributed neural circuits. Within the basal ganglia loop, interactions between distinct brain regions—particularly the primary motor cortex (M1) and the dorsal striatum (STR)—are central to motor planning, execution, and learning(1–5). Functional coupling between these regions facilitates integrated sensorimotor processing and adaptive behavior in healthy individuals.

Disruption of inter-regional functional connectivity, especially between M1 and the striatum, are hallmarks of several neurodegenerative movement disorders (6, 7). In Parkinson’s disease (PD), a neurodegeneration characterized by progressive motor impairment, functional magnetic resonance imaging (fMRI) studies consistently report reduced M1-STR connectivity(8–10). This cortico-striatal decoupling correlates with both the severity of motor symptoms and the duration of disease progression(11, 12), suggesting its potential as a biomarker for disease staging and treatment response. Similar patterns of disrupted connectivity are observed in other basal ganglia-related disorders, such as dystonia(13) and Huntington’s disease (14), further highlighting the importance of cortico-striatal interactions in motor control and dysfunction. Despite these insights, the mechanisms and onset of cortico-striatal decoupling in disease states remain poorly understood.

The striatum is primarily composed of GABAergic medium spiny neurons (MSNs), which account for over 90% of its neuronal population(15). These MSNs fall into two major subtypes: dopamine D1 receptor-expressing neurons (dMSNs), which participate in the direct pathway, and dopamine D2 receptor-expressing neurons (iMSNs), which contribute to the indirect pathway(16–19). Both subtypes receive converging excitatory input from the cortex, including projections from the primary motor cortex (M1), yet their respective contributions to motor control remain a subject of ongoing investigation and debate. Classical models propose that activation of dMSNs promotes movement through facilitation of thalamocortical activity, whereas iMSNs suppress movement via inhibition of thalamic output. However, emerging evidence challenges this dichotomy, suggesting more nuanced or overlapping roles during action initiation and suppression(20). The question of how these two MSN subtypes participate in cortico-striatal functional coupling during motor behavior remains largely unresolved.

In dopamine-depleted rodent models of PD, altered activity in both dMSNs and iMSNs has been reported(21). Some studies have shown similar reductions in firing rates or convergence of discharge patterns between the two populations following dopamine loss(22, 23). Conversely, other reports support the hypothesis that an imbalance between hyperactive iMSNs and hypoactive dMSNs is a critical driver of motor impairment(24–27). Despite these insights, it remains unclear whether such changes in MSN activity directly contribute to cortico-striatal decoupling and the progression of parkinsonian motor deficits. Dissecting the cell type-specific contributions of dMSNs and iMSNs to functional connectivity changes in PD is essential for advancing our understanding of disease mechanisms.

The intrastriatal injection of α-Syn preformed fibril (PFF) effectively induces aggregation of α-Syn and dopaminergic (DA) neuron cell loss in the substantia nigra (SN), two pathological hallmarks of PD(28) along with gradually advanced motor dysfunction(29–31). This model provides a valuable platform to examine whether cortico-striatal decoupling emerges prior to overt behavioral impairments.

In this study, we first established a dual-site fiber photometry system with cell type specificity to assess behavior-linked cortico-striatal coupling in vivo. Using the PFF-induced PD-like mouse model, we then applied this approach to analyze the temporal and cellular characteristics of cortico-striatal decoupling during disease progression. Our findings validate a fiber photometry-based method to study functional coupling with cell type resolution, and reveal the early emergence of behavior-specific decoupling between M1 and the dorsal striatum. Moreover, we demonstrate distinct contributions of dMSNs and iMSNs to this decoupling, offering new insight into the pathophysiological mechanisms underlying early-stage PD.

## RESULTS

### Dual-site fiber photometry enables monitoring of activity coupling between motor cortex and striatum during different motor behaviors

Several techniques, including electroencephalography (EEG), local field potential (LFP), and fMRI, have been widely used to assess neural coupling between brain subregions(32–34). However, these approaches are limited in their capacity to elucidate the underlying cellular mechanisms. Considering that striatal neurons are primarily composed of dMSNs and iMSNs, there is a pressing need for methods that enable cell type-specific analysis of cortico-striatal coupling during motor behavior in both healthy and diseased brains.

To address this challenge, we employed a dual-site fiber photometry approach capable of simultaneously monitoring calcium dynamics in genetically defined neuronal populations across multiple brain regions in freely behaving mice(35). Two calcium indicators, jRGECO1a (red) and GCaMp7s (green), were respectively expressed in neurons in the deeper layers of the primary motor cortex (M1) and dorsal striatum (STR) of the same hemisphere, followed by the implantation of two optical fibers (Fig. 1A-B; see Methods). This configuration enabled concurrent measurement of M1 and striatal activity without detectable spectral crosstalk (Fig. 1C, Fig. S1A).

**Figure 1.**
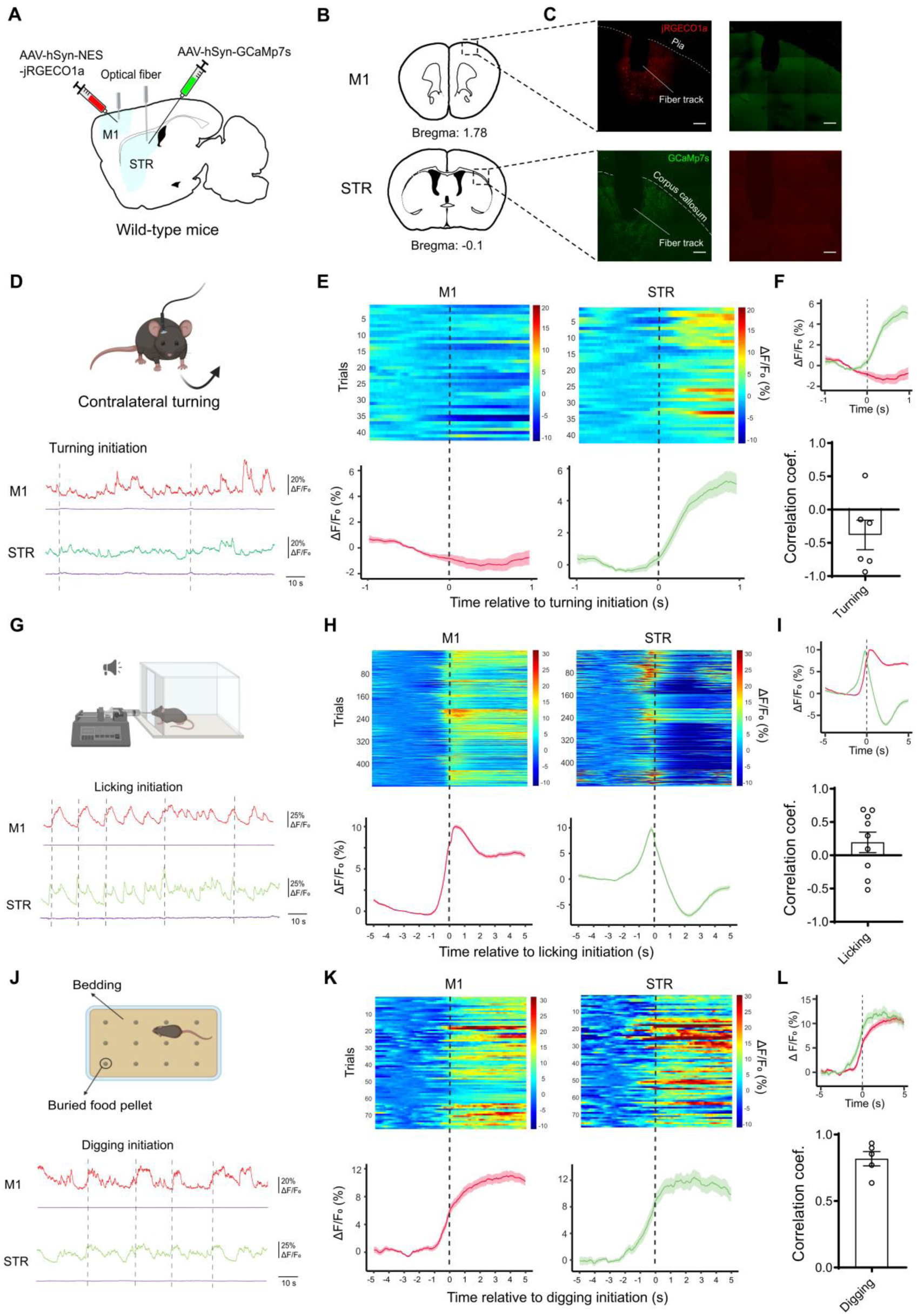
Detection of behavior-specific cortico-striatal coupling via dual-site fiber photometry imaging. **A.** Scheme of virus injection and optical fiber implantation. **B.** Locations (highlighted by dashed box) of virus injection and fiber placement. **C.** Anatomical verification of viral expression and optical fiber placement. Scale bar: 200 μm. **D-L**. Examination of the cortico-striatal coupling during contralateral turning (C-F), licking (G-I), and digging (J-L). (D, G, J) Behavioral diagrams (top) and example Ca^2+^ activities recorded from the primary motor cortex (M1, red) and the striatum (STR, green) with dotted lines denoted onsets of the specified behaviors. Ca^2+^-independent fluorescence was monitored using 410 nm light pulses for individual channel (purple, bottom). (E, H, K) Heatmaps showing all compiled trials (Top panels) and the averaged Ca^2+^ signal intensities (ΔF/F₀, bottom panels) aligned (dotted lines) to the initiation of the contralateral turning (D), licking (G), and digging (J). All left panels, M1 signal; all right panels, STR signal. (F, I, L) Combined M1 and striatal averaged Ca^2+^ activities temporally aligned to the initiation of behavior (Top panels) and calculated cortico-striatal correlation coefficient (Bottom panels) for contra-lateral turning (F), licking (I), and digging (L).

We focused on three ethologically relevant motor behaviors—contralateral turning, licking, and digging—to evaluate cortico-striatal coupling (Fig. 1D, 1G, 1J). All three behaviors reliably evoked neural activities with distinct dynamics in at least one brain regions (Fig. 1E, 1H, 1K, Fig. S1B-D). The degree of coupling was quantified by calculating Pearson’s correlation coefficients (hereafter referred to as “correlation coefficient”) between ΔF/F₀ calcium traces from M1 and STR within behavior-specific time windows (2 s for turning, 10 s for licking and digging; See Methods) centered around behavioral initiation. Licking behavior was automatically identified using a sensor, whereas contralateral turning and digging were manually annotated (see Methods). Consistent with previous findings(36), contralateral turning reliably evoked obvious striatal activation (Fig. 1E right, Fig S1B), validating the temporal precision of fiber photometry in monitoring fast motor behavior-associated neuronal activity. In contrast, M1 activity showed a modest suppression during turning (Fig. 1E left), resulting in a negative correlation coefficient between M1 and the striatum (Fig. 1F bottom).

During tone-cued licking, both M1 and STR were activated. However, the striatal signal exhibited a more phasic temporal profile relative to the broader M1 activation (Fig. 1G-I, Fig S1C), yielding an average correlation coefficient near zero (Fig. 1I bottom). To further characterize the temporal dynamics of interregional coupling, we computed time-resolved correlation coefficients using a 4-second sliding window (Fig. S1E_1_, see Methods). These analyses revealed transient and asynchronous patterns of M1–STR coordination during licking (Fig. S1E_2_). In contrast, digging—a goal-directed behavior in which mice retrieved buried food pellets (see Methods)—elicited strong and temporally aligned activation in both M1 and STR (Fig. 1J-L, Fig S1D), resulting in a high correlation coefficient approaching 1 (Fig. 1L bottom, Fig. S1E_3_). These behavior-associated signal dynamics were independent of calcium sensors used, as swapping the fluorescent sensors yielded comparable correlation values for licking and digging (Fig. S1F).

Together, these results demonstrate that our dual-site fiber photometry system effectively captures region- and behavior-specific neural dynamics. We identify distinct patterns of M1–STR coupling across motor behaviors, suggesting that cortico-striatal communication follows behavior-dependent coordination rules.

### dMSNs and iMSNs distinctively contribute to the cortico-striatal coupling in different motor behaviors

To dissect the contribution of different striatal MSNs to cortico-striatal coupling, GCaMP7s was selectively expressed in either striatal dMSNs or iMSNs by using D1R-Cre or D2R-Cre mouse lines, respectively (Fig. 2A-B; see Methods). Although striatal cholinergic interneurons (ChIs) also express D2Rs(37), they constituted only ∼1% of all GCaMP-positive cells in D2R-Cre mice (Fig. S2C-D), consistent with their known sparsity in the striatum(37). Thus, calcium signals recorded in D2R-Cre mice primarily reflected iMSN activity.

**Figure 2.**
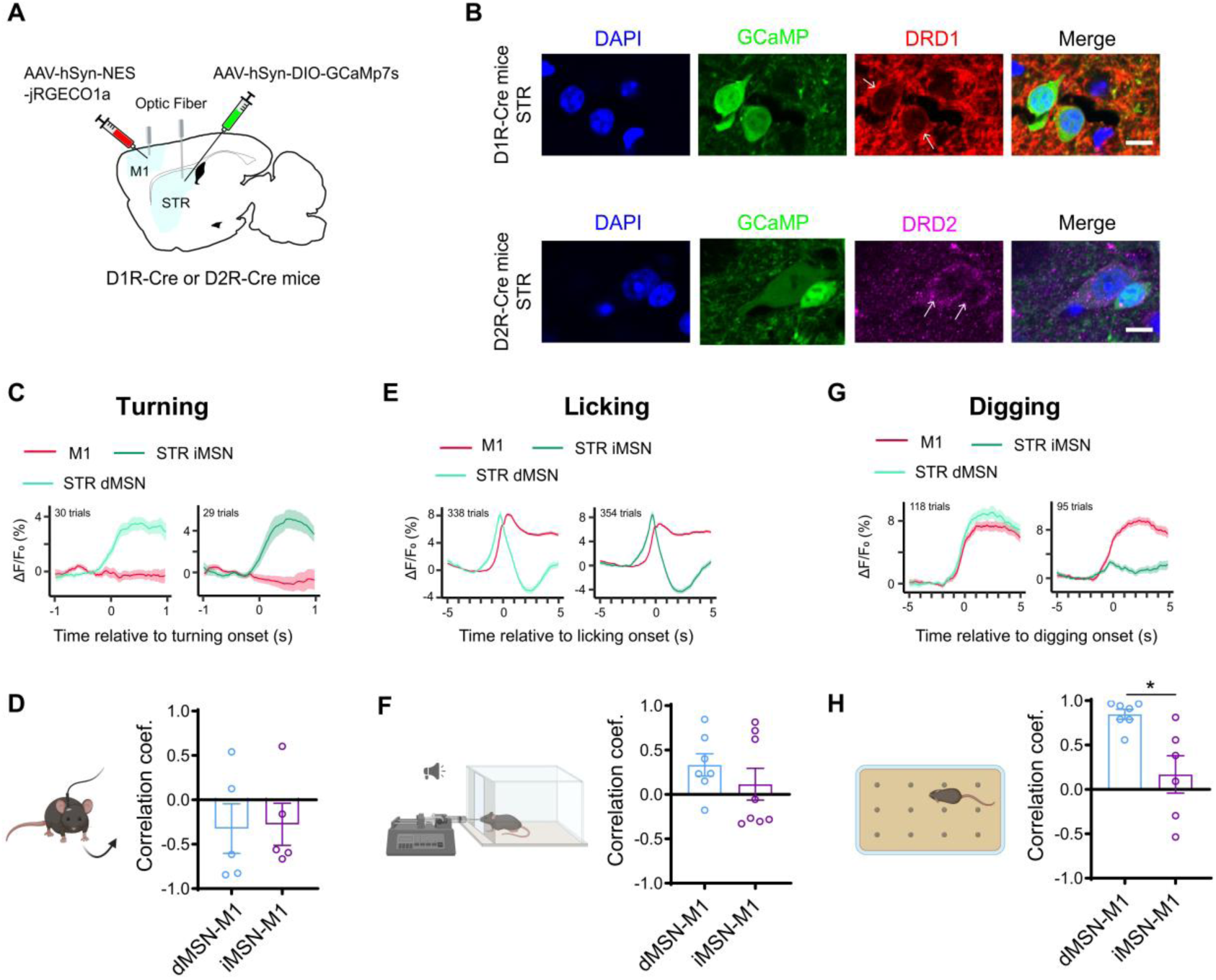
Behavior- and cell type-specific cortico-striatal coupling. **A.** Schematic diagram showing the experimental design for analyzing cell type-specific cortico-striatal coupling. **B.** Immunofluorescence images showing the colocalization between DRD1 (red) and DIO-GCaMP (green) in D1R-Cre mice (top panels), and the colocalization between DRD2 (magenta) and DIO-GCaMP (green) in the D2R-Cre mice (bottom panels). Scale bar: 10 μm. **C, E, G**. Average traces (ΔF/F₀) of M1 (red) and STR dMSNs (light green) and iMSNs (dark green) Ca^2+^ activities temporally aligned to the initiation of contralateral turning (C, dMSNs: 30 trials from 5 mice; iMSNs: 29 trials from 5 mice), licking (E, dMSNs: 338 trials from 7 mice; iMSNs: 354 trials from 8 mice), and digging (G, dMSNs: 118 trials from 7 mice; iMSNs: 95 trials from 6 mice). **D, F, H**. Comparison of cell type-specific cortico-striatal correlation coefficients for turning (D), licking (F), and digging behavior (H, *P* = 0.014). Mann-Whitney test.

Using dual-site fiber photometry, we simultaneously monitored M1 calcium activity and that of either dMSNs or iMSNs in the dorsal striatum (Fig. S2A-B). During contralateral turning and licking behaviors, both dMSNs and iMSNs exhibited comparable activity patterns (Fig. 2C-F) and similar coupling to M1 signals, as reflected by overlapping distributions of correlation coefficients (Fig. 2D, F). These observations are consistent with previous findings suggesting convergent involvement of both MSN subtypes in these behaviors (36, 38, 39).

Strikingly, digging behavior revealed a pronounced divergence in cortico-striatal coupling between the two cell types. dMSNs showed strong activation resembling the overall striatal activity observed in wild-type mice and exhibited robust temporal coupling with M1 signals (Fig. 2G-H). In contrast, although iMSNs were active during general locomotion, turning, and licking (Fig. S2B, Fig. 2C–F), their responses during digging were markedly attenuated (Fig. 2G, Fig. S2F), resulting in a significantly weaker M1– iMSN coupling compared to that of M1–dMSN (Fig. 2H, Fig S2E_2_).

These findings highlight the cell type-specific nature of M1-striatal coupling during motor behaviors and underscore the dominant role of dMSNs in digging-associated activity.

### Intrastriatal injection of α-syn PFF induces PD-like pathology in wild-type mice

Cortico-striatal decoupling is a well-documented feature of PD, observed across mild to severe stages and correlating with motor impairments(11, 40, 41). To investigate the onset and mechanism underlying this PD-associated cortico-striatal decoupling, we adapted an inducible PD-like mouse model established previously(42) by bilaterally inoculating α-synuclein (α-Syn) preformed fibrils (PFF) into the dorsal striatum of wild-type mice (Fig. 3A). Behavioral and pathological phenotypes were examined at both one and three months post PFF inoculation (Fig 3B).

**Figure 3.**
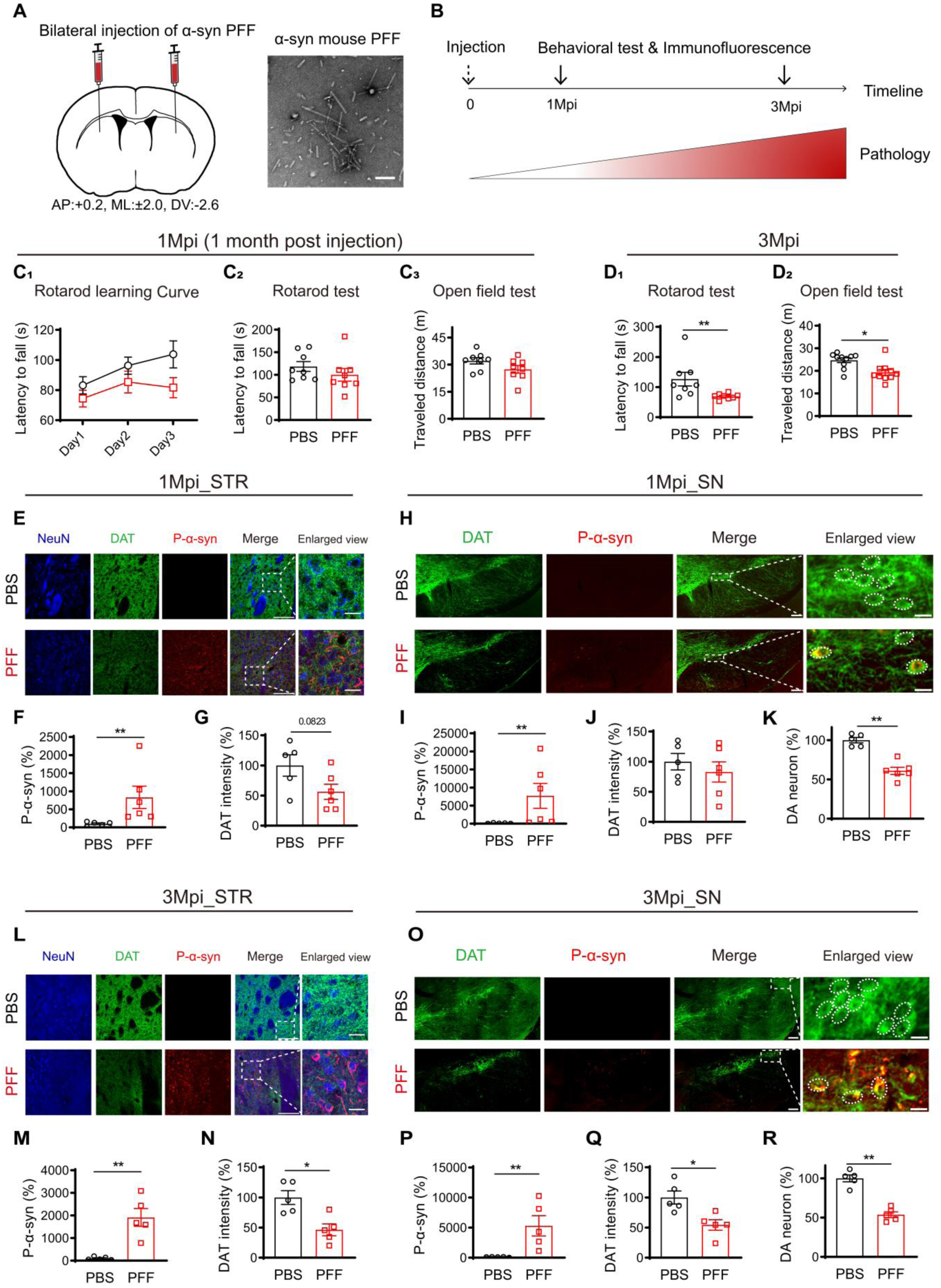
Pathological characterization of α-synuclein preformed fibril (PFF) induced PD-like mouse model. **A.** Schematic diagram describing the bilateral inoculation of PFF into striatum of wild-type mice. Right panel, transmitted electron microscope image of PFF. Scale bar: 100 nm. **B.** Experimental timeline. **C-D.** Behavioral analysis of the PFF- and PBS-injected mice at 1 (C) and 3 (D) months post injection (Mpi). (C_1_) Learning curve of rotarod running (Two-way ANOVA, F (_1, 14_) = 2.868, *P* = 0.1125). (C_2_) Latency to fall in the rotarod test at 1 Mpi. (C_3_) Total travel distance in the open field test at 1 Mpi. (D_1_) Latency to fall in the rotarod test at 3 Mpi (*P* = 0.0207). (D_2_) Total travel distance in the open field test at 3 Mpi (*P* = 0.0147). **E-K.** Immunofluorescence analyses of pathological hallmarks in PFF- and PBS-injected mice at 1 Mpi. All data normalized to PBS control. NeuN, neuron-specific nuclear protein. (E-G) Comparison of the p-α-Syn intensity (F, *P* = 0.0043) and dopamine transporter (DAT) signal intensities (G) in STR. E, Representative immunofluorescence images. Scale bars: 100 μm; 20 μm for enlarged view. (H-K) Comparison of the p-α-Syn intensity (I, *P* = 0.0043), DA neuron number (J, *P* = 0.0043) and DAT intensity (K) in substantia nigra (SN). H, Representative immunofluorescence images. Scale bars: 100 μm; 20 μm for enlarged view. **L-R.** Immunofluorescence analyses of pathological hallmarks in PFF- and PBS-injected mice at 3 Mpi. All data normalized to PBS control. (L-N) Comparison of the p-α-Syn intensity (M, *P* = 0.0079) and dopamine transporter (DAT) signal intensities (N, *P* = 0.0317) in STR. L, Representative immunofluorescence images. Scale bars: 100 μm; 20 μm for enlarged view. (O-R) Comparison of the p-α-Syn intensity (P, *P* = 0.0079), DA neuron number (Q, *P* = 0.0079) and DAT intensity (R) in SN. O, Representative immunofluorescence images. Scale bars: 100 μm; 20 μm for enlarged view. Mann-Whitney test.

At one-month post-injection (1 Mpi), PFF-treated mice showed a mild reduction in rotarod learning ability (Fig. 3C_1_) without impairments in rotarod test performance (Fig. 3C_2_) or general locomotor activity (Fig. 3C_3_). By three months post injection (3 Mpi), mice exhibited pronounced motor deficits (Fig. 3D_1_, D_2_), reflecting progressive PD-like motor impairments. We further assessed two key pathological markers of PD: α-Syn aggregates and dopaminergic neuron loss, using immunostaining. PFF inoculation led to a significant, time-dependent increase in phosphorylated α-Syn (p-α-Syn) immunoreactivity, a hallmark of synucleinopathy, in both the striatum (Fig. 3E-F, L-M) and the substantia nigra (SN) (Fig. 3H-I, O-P) at 1 and 3 Mpi. The rapid formation and propagation of synucleinopathy via direct synaptic connectivity(43) coincided with a reduction in striatal dopamine transporter (DAT) signal intensity (Fig. 3Q) and a decreased dopaminergic neuronal cell count in the SN beginning at 1 Mpi (Fig. 3K, R). In contrast, the ventral tegmental area (VTA), which is mainly innervated by MSNs in the ventral striatum, exhibited milder phosphorylated α-synuclein (p-α-Syn) immunoreactivity and delayed onset of dopaminergic neuron loss (Fig. S3A-H).

These findings confirm that PFF inoculation induces α-Syn aggregate formation, which spreads to the SN and drives the progressive pathology of dopaminergic neurons in a time-dependent manner.

### Progressive and behavior-specific cortico-striatal decoupling in the PFF-injected mice

Using the PFF-induced PD-like mouse model, we investigated the onset and mechanisms of cortico-striatal decoupling during disease progression via dual-site fiber photometry. PFF inoculation did not affect fiber-based calcium imaging quality (Fig. S4A-C). Given its strong coupling, we compared digging-associated M1-dorsal striatum coupling between PFF- and PBS-injected mice. At one-month post-injection (1 Mpi), M1 and striatal responses during digging were largely preserved in PFF mice (Fig. 4C, D_3_), and the M1–striatum correlation coefficient did not differ significantly from controls (Fig. 4D_1_). Striatal responses showed a trend of earlier onset compared to M1 responses at 1 Mpi (Fig. 4C), which became more obvious by 2 and 3 Mpi (Fig 4E & G). To gain more detail temporal analysis, we computed the phase lag—the time shift required for maximal cross-correlation between M1 and striatal signals—as a metric of response synchrony (Fig. 4B, see Methods). While a slight increase in phase lag was observed in PFF mice at 1 Mpi (Fig. 4D_2_), this difference became statistically significant by 2 Mpi (Fig. 4E, F_2_), accompanied by a decline in the cortico-striatal correlation coefficient (Fig. 4F_1_). Importantly, the amplitude of cortical and striatal calcium signals remained unchanged at this stage (Fig. 4F_3_, Fig. S4E), indicating that decoupling occurred independently of gross activity levels.

**Figure 4.**
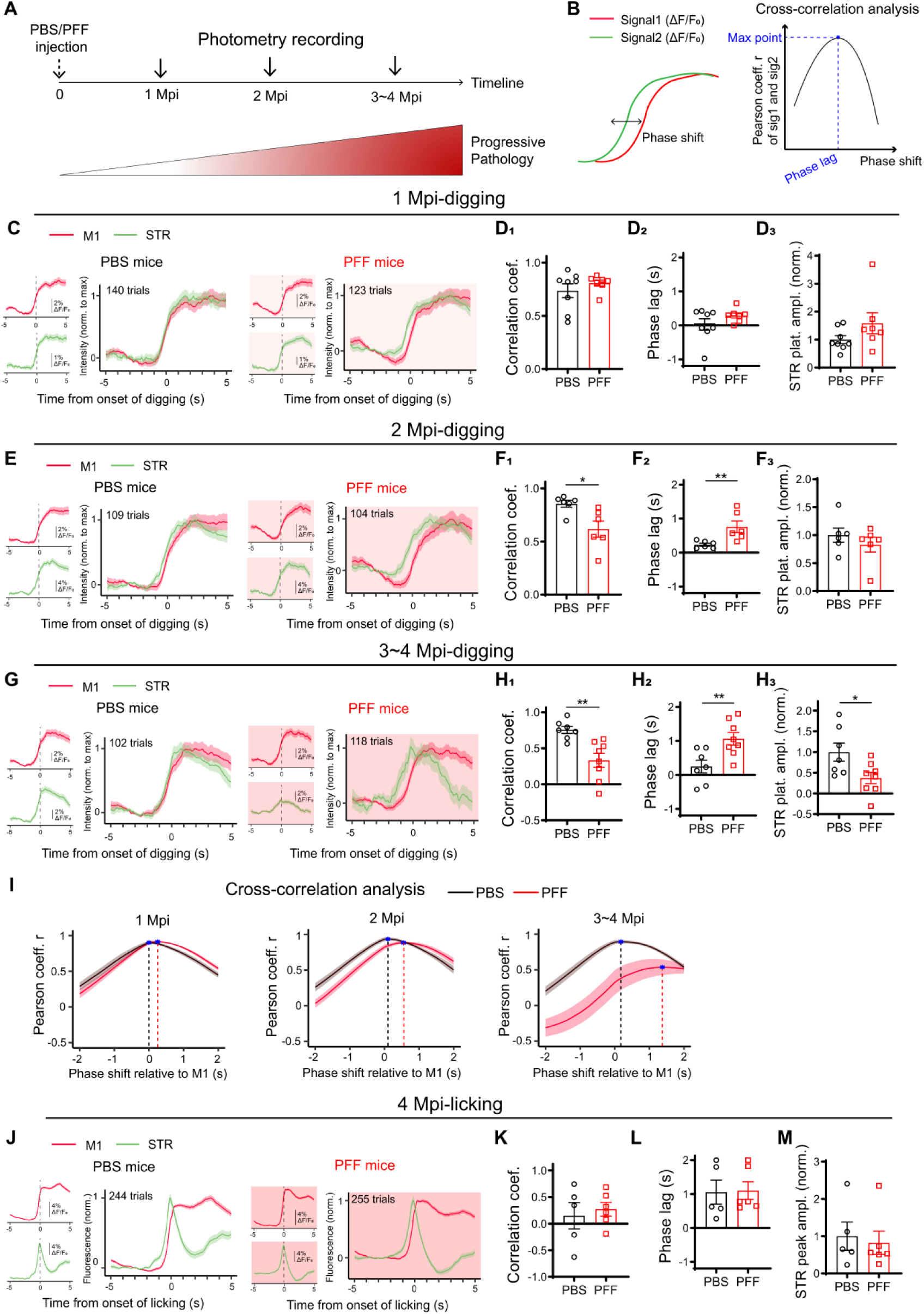
Behavior-specific cortico-striatal decoupling in PD-like mice. **A.** Experimental timeline. **B.** Schematic diagram illustrating how the phase lag was calculated. **C-I,** Dual-site fiber photometry imaging of M1 and STR in PBS- and PFF-injected mice at 1 Mpi (C-D), 2 Mpi (E-F), and 3-4 Mpi (G-H) during digging. (C, E, G) ΔF/F₀ traces of M1 (red) and STR (green) Ca^2+^ activities temporally aligned to the initiation of digging in mice injected with either PBS (panels with white background) or PFF (panels in pink background). The average ΔF/F₀ waveform of M1 (Top left panels) and STR (Bottom left panels) were presented, and the waveform for both regions were overlaid after normalizing each channel to its maximum value (panels on the right). (D, F, H) Quantitative analysis of digging-associated cortico-striatal coupling and normalized striatal response amplitudes in PBS-injected (black) and PFF-injected (red) mice. The striatal response amplitudes were normalized to the PBS control values within each replicate. (D_1_, F_1_, H_1_) Correlation coefficient. (F_1_, *P* = 0.0152; H_1_, *P* = 0.0012). (D_2_, F_2_, H_2_) Phase lags (F_2_, *P* = 0.0065; H_2_, *P* = 0.0051). (D_3_, F_3_, H_3_) Plateau amplitudes of striatal response. Data batch-normalized to the PBS group. (H_3_, *P* = 0.0401) (I) Mean cross-correlation between digging-associated M1 and STR signals from PBS-(black) and PFF-injected (red) mice at different time-points post injection. The x-coordinates of each peak highlighted by blue asterisks corresponded to the phase lag for each group. **J-M**. Dual-site fiber photometry imaging of M1 and STR at 4 Mpi during licking. (J) The average ΔF/F₀ waveform of M1 (Top left panel) and STR (Bottom left panel) and the overlaid normalized waveform (Right panel) temporally aligned to the initiation of licking in mice injected with PBS (panels with white backgrounds) and PFF (panels with pink backgrounds). (K) Comparison of the coupling indices. (L) Comparison of the phase lags. (M) Comparison of the licking-associated peak amplitudes of striatal response. Mann-Whitney test.

By 3–4 Mpi, PFF mice exhibited further deterioration in cortico-striatal coupling during digging. In contrast, PBS-injected controls maintained strong coupling throughout the same period (Fig. 4G, H_1_, Fig. S3K). This progressive decoupling in PFF mice was accompanied by a continued increase in phase lag (Fig. 4H_2_) and a significant reduction in striatal response amplitude (Fig. 4H_3_). Full-time-window cross-correlation analyses further demonstrated an early impairment in the temporal precision of cortico-striatal coordination (Fig. 4I), with progressive exacerbation as pathology advanced.

Importantly, this decoupling was behavior-specific Despite pronounced deficits in digging-associated M1–striatum coupling, PFF-injected mice retained normal coupling during licking behavior, even at 4 Mpi (Fig. 4J-M). These findings suggest that early PD-associated decoupling is behavior-specific.

### Disruption of M1–striatum coupling impairs digging behavior

We next examined whether cortico-striatal decoupling during digging behavior was associated with observable behavioral deficits. Digging behavior was quantified in terms of duration and frequency (see Methods). At 3–4 months post-injection (Mpi), PFF mice exhibited reduced digging duration (Fig. 5A) and increased digging frequency (Fig. 5B) compared to PBS controls.

**Figure 5.**
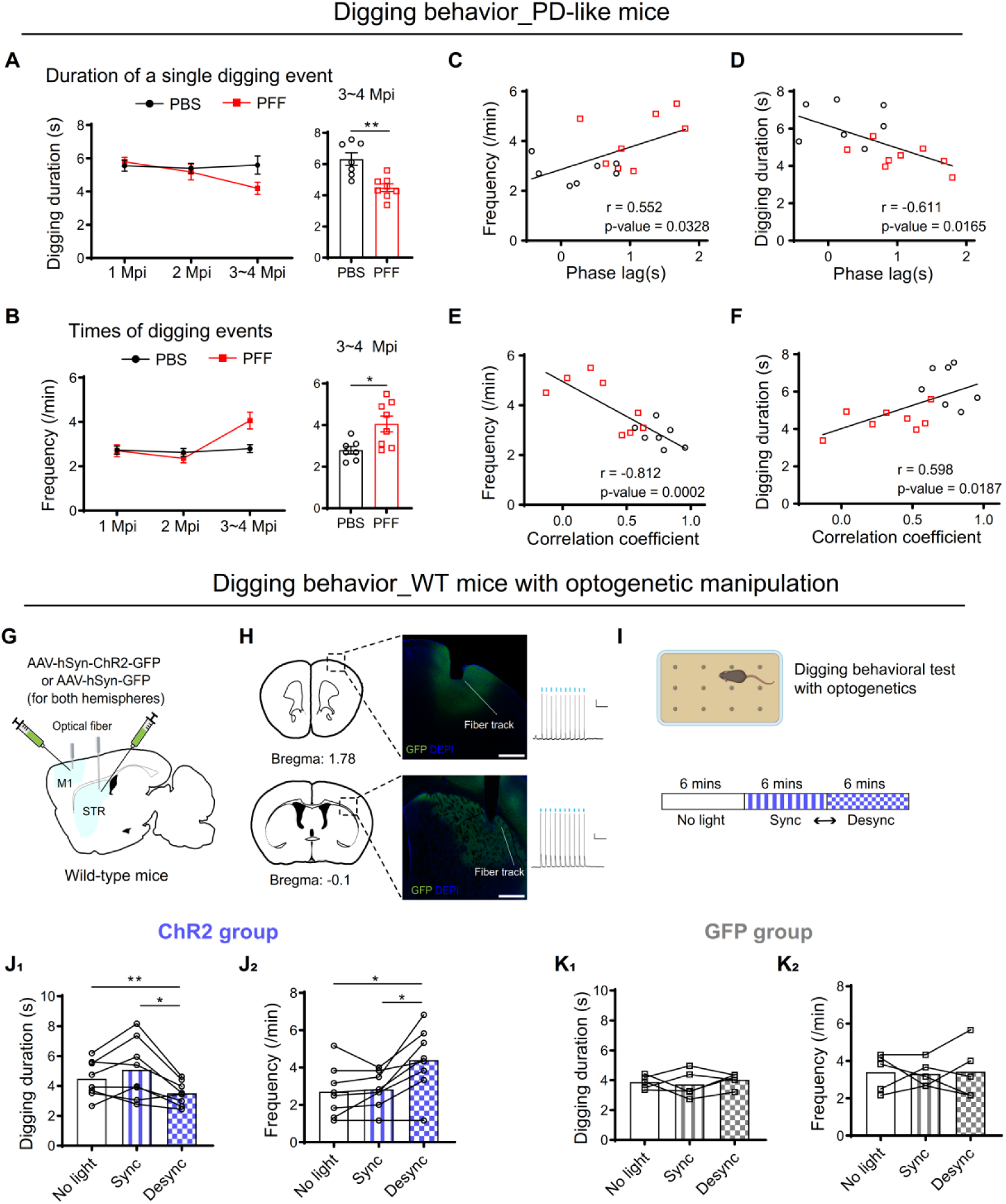
Cortico-striatal decoupling both correlates with and drives digging behavior deficits. **A.** Duration of a single digging event. Left, the summary of average digging duration across different time-points post injection. Right, comparison of the duration from mice injected with PBS and PFF at 3∼4 Mpi (Mann-Whitney test, *P* = 0.0022). **B.** Frequencies of digging event. Left, the summary of digging frequency across different time-points post injection. Right, comparison of the frequencies at 3∼4 Mpi (Mann-Whitney test, *P* = 0.0152) **C, D**. Correlation analysis of phase lags with digging frequencies (C) and duration (D) from mice injected with PBS (black) and PFF (red) mice at 3∼4 Mpi. r: Pearson’s correlation coefficient. Each dot represents data from one mouse (the same below). **E, F**. Correlation analysis of correlation coefficient with digging frequency (E) and duration (F) from mice injected with PBS (black) and PFF (red) at 3∼4 Mpi. **G.** Scheme of virus injection and optic fiber implantation for optogenetic stimulation. **H.** Left, anatomical verification of viral expression and optical fiber placement. Scale bar: 0.5 mm. Right, *ex vivo* electrophysiological verification confirmed that ChR2-expressing neurons reliably responded to blue light stimulation. **I.** Diagrams depicting the optogenetic stimulation protocols: three 6-minute epochs for both ChR2- and GFP-expressing mice: no light stimulation, synchronized stimulation of M1 and STR (Sync), desynchronized stimulation of M1 and STR (Desync). The order of the Sync and Desync epochs was randomly and evenly assigned. **J.** Effects of optogenetic stimulation on digging duration (J_1_, F_(1.325, 9.272)_ = 8.837, *P* = 0.0113; No light vs. Desync, *P* = 0.0084; Sync vs. Desync, *P* = 0.0368) or total digging events per epoch (J_2_, F _(1.261, 8.825)_ = 11.35, *P* = 0.0065; No light vs. Desync, *P* = 0.0395; Sync vs. Desync, *P* = 0.0140) in the ChR2 group. **K.** Effects of optogenetic stimulation on digging duration (K_1_, F _(1.464, 5.854)_ = 0.3714, *P* = 0.6437) or total digging times per epoch (K_2_, F _(1.979, 7.914)_ = 0.04056, *P* = 0.9594) in the GFP group. One-way ANOVA followed by Šídák’s multiple comparisons test.

To assess the relationship between cortico-striatal coordination and digging behavior, we analyzed the correlation between behavioral metrics and neural coupling parameters. Phase lag and correlation coefficient between M1 and striatum were significantly correlated with both digging frequency (Fig. 5C, E) and duration (Fig. 5D, F). In contrast, the amplitude of striatal calcium signals did not correlate with digging performance (Fig. S5C–D), indicating that the timing and coordination of activity, rather than activity magnitude, were critical for behavioral output. Moreover, altered digging patterns in PFF mice were associated with impaired task performance, as evidenced by a reduced success rate in food pellet retrieval (Fig. S5A–B). These findings suggest a strong link between M1–striatum functional coupling and effective execution of complex motor behavior. In contrast, M1–striatum coupling during licking remained intact in PFF mice, and their licking behavior was unaffected (Fig. S5E–H), further supporting a behavior-specific role of cortico-striatal coordination.

Given these observations and a prior study showing that disruption of inter-regional coupling via optogenetics can modulate behavior in mice (44), we sought to determine whether artificial disruption of M1–striatum coordination could causally impair digging. To this end, wild-type (WT) mice were bilaterally injected with AAVs expressing either Channelrhodopsin-2 (ChR2) or GFP in M1 and the dorsal striatum, followed by optical fiber implantation (Fig. 5G, see Methods). Whole-cell recordings from acute brain slices confirmed that blue light pulse stimulation effectively activated neurons expressing ChR2 (Fig. 5H).

Four weeks post-surgery, mice were subjected to three digging test conditions: 6 minutes of digging without light stimulation (No Light), followed by 6 minutes each of synchronized (Sync) or desynchronized (Desync; 90° out-of-phase) bilateral optogenetic stimulation of M1 and striatum. The Sync and Desync conditions were applied in randomized order (Fig. 5I). Light stimulation of M1 or striatum alone had no impact on digging behavior (Fig. S5I-J), nor did synchronized stimulation of both regions (Fig. 5J, middle bars). However, desynchronized stimulation significantly reduced digging duration and increased digging frequency in ChR2-expressing mice (Fig. 5J, right bars), mimicking the behavioral phenotype observed in PFF mice. These effects were absent in GFP-expressing control animals (Fig. 5K).

Taken together, these findings demonstrate that precise temporal coordination between M1 and striatal activity is critical for executing complex motor behaviors. Disruption of this coordination—either through pathology or artificial desynchronization—leads to behaviorally relevant deficits, highlighting the importance of functional connectivity rather than isolated activity within individual brain regions.

### Defects in dMSNs neuronal activity mediates the digging-associated cortico-striatal decoupling in PD-like mice

A functional imbalance between the dMSNs and iMSNs has been proposed to underlie the bradykinesia in PD patients(45). To determine whether cell-type-specific alterations contribute to the early-onset cortico-striatal decoupling observed in PFF-induced PD-like mice, we employed a strategy similar to that described in Figure 2A (Fig. 6A). D2R-Cre mice with CreOFF-GCaMP expression in the striatum (See Methods) could effectively label dMSNs (Fig. S6A) and generate reliable digging-associated striatal responses (Fig S6B) resembled those observed in D1R-cre mice expressing CreON-GCaMP (Fig. S6C). Therefore, M1-dMSNs coupling in the context of PD development was analyzed in both D1R-cre mice with CreON-GCaMP (Fig. 6B-E, Open symbols) and D2R-Cre mice with CreOFF-GCaMP (Fig. 6B-E, Filled symbols).

**Figure 6.**
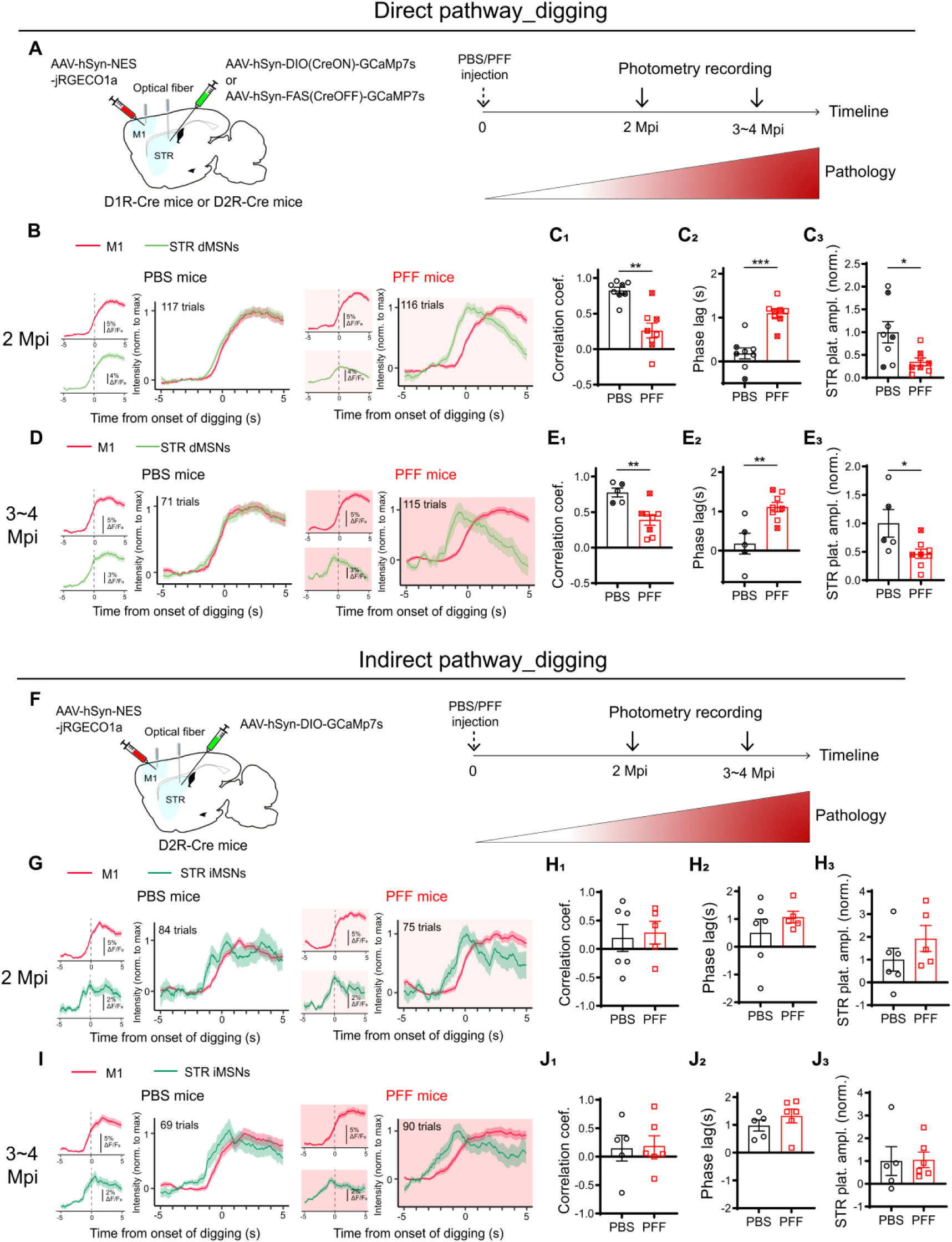
Cell type-specific contribution of digging-related cortico-striatal decoupling in PFF-injected mice. **A-E.** Dual-site fiber photometry imaging of M1 and striatal dMSNs in PFF- and PBS-injected mice at 2 Mpi (B-C), and 3-4 Mpi (D-E) during digging. (A) Schematic illustration of the viral injection and optical fiber implantation strategy for monitoring M1-dMSNs coupling. A Cre-dependent GCaMP virus (CreON-GCaMP) was injected into the dorsal striatum of D1R-Cre mice to label dMSNs, while a Cre-suppressible GCaMP virus (CreOFF-GCaMP) was used in D2R-Cre mice to target iMSNs. Right panel, experiment timeline. (B, D) ΔF/F₀ traces of M1 (red) and striatal dMSNs (light green) Ca^2+^ activities temporally aligned to the initiation of digging at 2 Mpi (B) and 3∼4 Mpi (D) in mice injected with either PBS (panels with white background) or PFF (panels in pink background). The average ΔF/F₀ waveform of M1 (Top left panels) and STR (Bottom left panels) were presented, and the waveform for both regions were overlaid after normalizing each channel to its maximum value (panels on the right). (C, E) Quantitative analyses of the M1-dMSNs coupling and normalized plateau amplitudes of dMSNs response at 2 Mpi (C) and 3-4 Mpi (E). Open symbols were data from CreON-GCaMP mice; the cross-filled symbols were data from CreOFF-GCaMP mice. (C_1_, E_1_) Correlation coefficients. (C_1_, *P* = 0.0011. E_1_, *P* = 0.0062). (C_2_, E_2_) Phase lags (E_2_, *P* = 0.0003. G_2_, *P* = 0.0062). (C_3_, E_3_) Plateau amplitudes of dMSNs Ca^2+^ activities (C_3_, *P* = 0.0148. E_3_, *P* = 0.0451). Data normalized to the PBS group. **F-J.** Dual-site fiber photometry imaging of M1 and striatal iMSNs in PFF- and PBS-injected mice at 2 Mpi (G-H), and 3-4 Mpi (I-J) during digging. Right panel, experiment timeline. (F) Scheme of virus injection and optical fiber implantation. (G, I) Same as in B and D but for M1 (red) and striatal iMSNs (dark green) Ca^2+^ activities at 2 Mpi (G) and 3∼4 Mpi (I). (H, J) Quantitative analyses of the M1-iMSNs coupling and normalized plateau amplitudes of iMSNs Ca^2+^ activities at 2 Mpi (H) and 3-4 Mpi (J). (H_1_, J_1_) Comparison of the correlation coefficients. (H_2_, J_2_) Comparison of the phase lags. (H_3_, J_3_) Comparison of the plateau amplitudes of iMSNs Ca^2+^ activities. Data normalized to the PBS group. Mann-Whitney test.

To analyze M1–dMSN coupling during disease progression, we recorded from both D1R-Cre mice (CreON-GCaMP; Fig. 6B–E, open symbols) and D2R-Cre mice (CreOFF-GCaMP; Fig. 6B–E, filled symbols). At 2 months post-injection (2 Mpi), when overall cortico-striatal decoupling begins to manifest in PFF mice, digging-associated responses in dMSNs were already significantly decoupled from M1 activity (Fig. 6B). This was evidenced by a significant reduction in correlation coefficient (Fig. 6C_1_) and increased phase lag (Fig. 6C_2_). Notably, the amplitude of dMSN calcium responses was also significantly reduced (Fig. 6C_3_), preceding the decline in overall striatal activity observed at later stages (see Fig. 4H_3_).

By 3–4 Mpi, M1–dMSN decoupling further worsened (Fig. 6D–E), whereas coupling remained largely intact at 1 Mpi (Fig. S7F-G), indicating a progressive, cell type-specific disruption. In contrast, M1–iMSN coupling during digging was preserved throughout the disease course in PFF-injected mice (Fig. 6F– J), suggesting that decoupling was specific to the direct pathway.

Importantly, PFF inoculation did not alter M1–dMSN or M1–iMSN coupling during licking behavior (Fig. S7H–I), indicating that the observed decoupling is both cell type- and behavior-specific in this PD-like model.

### Activation of D1R but not D2R rescues the cortico-striatal decoupling and behavior defects of the PFF-injected mice

Insufficient striatal dopamine release from substantia nigra (SN) dopaminergic terminals is the primary driver of basal ganglia circuit dysfunction underlying motor deficits in PD(45). Recent studies have linked dopamine depletion to abnormal functional connectivity in both rodent models and PD patients(46–49). Consistent with this, PFF-injected mice exhibited reduced striatal dopamine levels during locomotion (Fig. S8F-H).

To investigate whether the impaired dopamine release is responsible for the dMSN-specific cortico-striatal decoupling and related behavior deficit, PFF mice at 4-month post-injection were administered intraperitoneal (i.p.) injections of L-DOPA, SKF-81297 (SKF, a D1R receptor agonist), quinpirole (Quin, a D2R receptor agonist), or saline (SAL) (Fig. 7A). None of the treatments altered M1 calcium responses during digging (Fig. S8A-B). Consistent with earlier findings, both L-DOPA and SKF—but not Quin— reduced phase lag between M1 and striatal activity (Fig. 7C) and significantly increased cortico-striatal correlation (Fig. 7D), suggesting restored coupling. Additionally, both L-DOPA and SKF elevated striatal response amplitude, while Quin had no such effect (Fig. 7E). These results support the conclusion that dopamine depletion disrupts cortico-striatal communication by impairing dMSN activation during digging.

**Figure 7.**
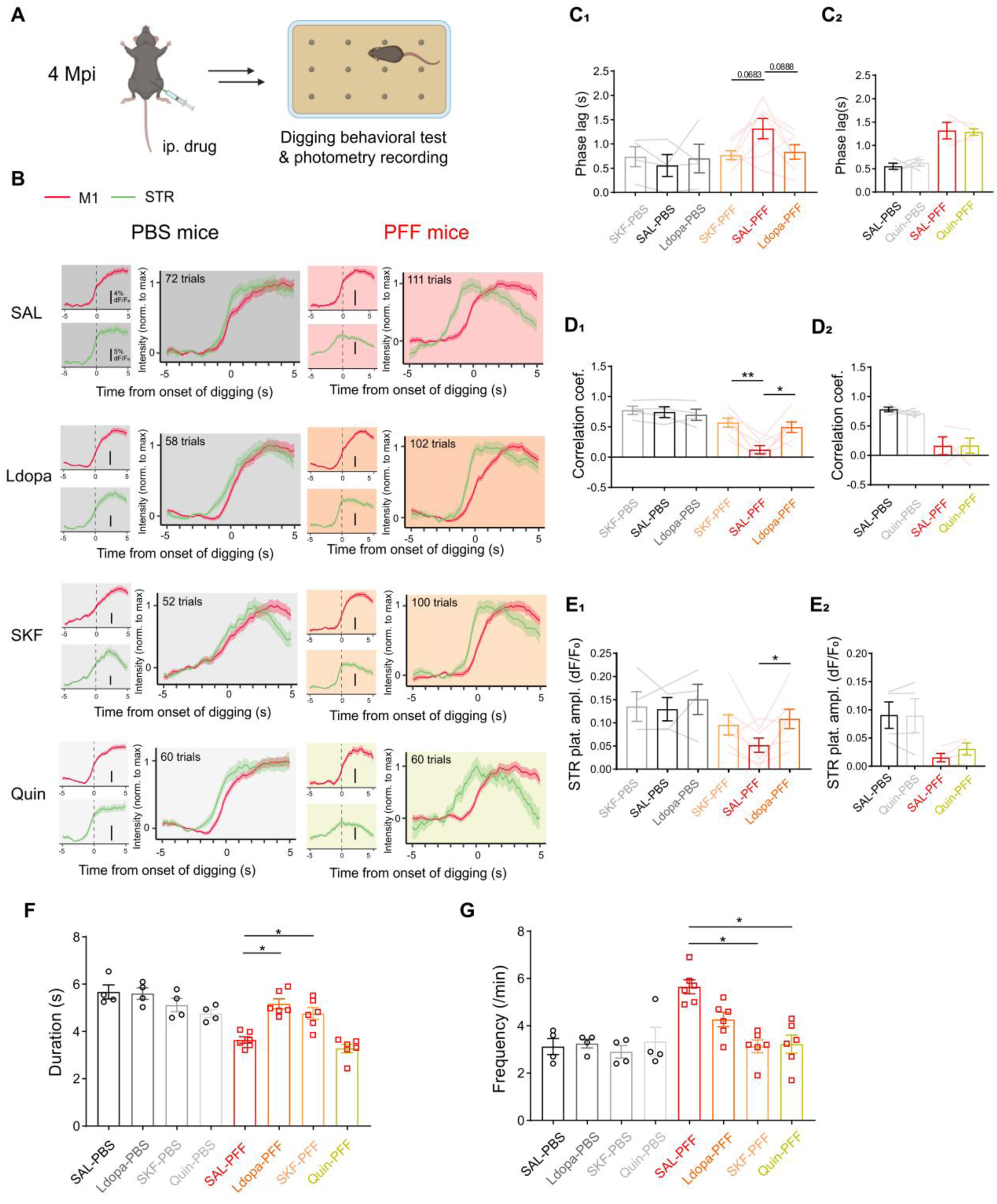
Selective activation to D1R rescues both digging-associated cortico-striatal decoupling and digging behavioral defects. **A**. Schematic diagram for pharmacological experiments. i.p., intraperitoneal administration. **B-E.** Dual-site fiber photometry imaging of M1 and STR in PFF- and PBS-injected mice at 4 Mpi during digging, after ip. SAL (saline), L-dopa, SKF (SKF-81297, a D1R selective agonist), or Quin (quinpirole, a D2R selective agonist). (B) ΔF/F₀ traces of M1 (red) and STR (green) Ca^2+^ activities temporally aligned to the initiation of digging in mice injected with either PBS (panels with white background) or PFF (panels in pink background) after administration of drugs. The average ΔF/F₀ waveforms of M1 (Top left panels, scale bars: 4% ΔF/F₀) and STR (Bottom left panels, scale bars: 5% ΔF/F₀) were presented, and the waveforms for both regions were overlaid after normalizing each channel to its maximum value (panels on the right). (C, D, E) Quantitative analyses of the digging-associated cortico-striatal coupling (C, D) and amplitudes of striatal Ca^2+^ activities (E) after administration of drugs. (C_1_) Comparison of the phase lags (PFF groups: SAL-PFF as control, F_(1.821, 10.93)_ = 4.960, SAL-PFF *vs.* L-dopa-PFF, *P* = 0.0888; SAL-PFF vs. SKF-PFF, *P* = 0.0683). (C_2_) Comparison of the phase lags after saline or quinpirole administration. (D_1_) Comparison of the correlation coefficient (PFF groups: SAL-PFF as control, F_(1.561, 9.637)_ = 11.92, SAL-PFF *vs.* L-dopa-PFF, *P* = 0.0314; SAL-PFF vs. SKF-PFF, *P* = 0.0011). (D_2_) Comparison of the correlation coefficient after saline or quinpirole administration. (E_1_) Comparison of the plateau amplitudes of striatal activity (PFF groups: SAL-PFF as control, F_(1.453, 8.719)_ = 6.152, SAL-PFF vs. L-dopa-PFF, *P* = 0.0459). (E_2_) Comparison of the plateau amplitudes of striatal activity, after saline or quinpirole administration. **F-G.** Effect of dopaminergic drugs on digging behavior. (F) Comparison of digging duration (PFF groups: SAL-PFF as control, F_(1.463, 7.317)_ = 19.97, SAL-PFF *vs.* Ldopa-PFF, *P* = 0.0018; SAL-PFF *vs.* SKF-PFF, *P* = 0.0388). (G) Comparison of digging frequency (PFF groups: SAL-PFF as control, F_(2.120, 10.60)_ = 11.78, SAL-PFF *vs.* SKF-PFF, *P* = 0.0143; SAL-PFF *vs.* Quin-PFF, *P* = 0.0192). One-way ANOVA followed by Dunnett’s multiple comparisons test, unless otherwise noted. Wilcoxon matched-pairs test (C_2_, D_2_, E_2_).

We next assessed the behavioral impact of these pharmacological interventions. Both L-DOPA and SKF significantly improved digging performance in PFF mice (Fig. 7F–G), without affecting general locomotion (Fig. S8C). Notably, SKF treatment also increased task success rates, further implicating dMSN-specific pathways in rescuing behavioral deficits (Fig. S8D). In contrast, Quin treatment failed to improve digging behavior (Fig. 7F, Fig. S8D), consistent with the preserved M1–iMSN coupling observed in PFF mice.

## DISCUSSION

Coordinated activity between the cortex and striatum is essential for the execution and refinement of motor behaviors, particularly skilled and goal-directed movements(4, 5). While several methods—such as LFP, EEG, and fMRI—have been developed to assess inter-regional brain coordination, they lack cell type-specific resolution(50). In this study, we utilized dual-site fiber photometry with genetically defined cell-type specificity to monitor dynamic, behavior-linked M1–striatal coordination in vivo. Although photometry offers lower temporal resolution than LFP or EEG(51, 52), it enables the detection of M1–striatum coupling patterns specific to distinct motor behaviors (Fig 1). Among the behaviors examined, spontaneous contralateral turning, considered a reflexive behavior(53), evoked moderate striatal activity but minimal M1 response, resulting in negative coupling. In contrast, both tone-cued licking and food pellet-driven digging, as goal-directed movements, induced robust activation in both regions but exhibited different correlation coefficients, reflecting distinct kinetics of calcium responses. These signal dynamics suggest differential regional engagement during motor behaviors. For example, Ca²⁺ signals in both M1 and striatum precede licking onset detected by the sensor, potentially representing motor planning(54). Moreover, the rapid decay of striatal activity post-initiation suggests that striatal neurons may play a role in action initiation, whereas M1 may contribute to both planning and execution (55). These findings demonstrate the utility of fiber photometry in elucidating functional brain coordination.

The roles of dMSNs and iMSNs in cortico-striatal coupling remain incompletely understood. Our results provide direct evidence that these neuronal subtypes contribute differentially to motor behavior-specific coordination. Consistent with previous findings(38, 56), both dMSNs and iMSNs were activated during turning and licking, contributing similarly to cortico-striatal coupling. However, during digging, M1 showed stronger coupling with dMSNs than iMSNs, aligning with earlier studies showing differential recruitment of MSN subtypes during tasks like lever pressing(18, 57). These findings emphasize the behavior-dependent contributions of dMSNs and iMSNs in shaping cortico-striatal dynamics.

The impaired digging behavior in PFF mice likely reflects deficits in forelimb function, a core feature of bradykinesia in PD animal models(58, 59) and patients(60). Progressive hesitation and shortened movement duration, hallmarks of bradykinesia(61), shares some similarity with the decreased digging duration and increased frequency in PFF mice. Importantly, M1–striatum coupling during digging, but not other behaviors, exhibited early disruption prior to observable behavior deficit. Given the asymmetric activity between dMSNs and iMSNs during digging, such imbalance nature might render digging more vulnerable to early PD pathology. Furthermore, forelimb hypokinesia has been shown linearly related to functional disconnection of striatal dMSNs and motor cortex neurons in another PD-like mouse model(62), which is consistent with our discovery.

Striatal fast-spiking interneurons (FSIs), known to regulate the timing and synchronization(63–65), may contribute to this vulnerability. FSIs provide lateral inhibition within striatal microcircuits(66) and exhibit faster activation and lower thresholds than MSNs(67, 68), making them key regulators of striatal dynamics. Their activity is modulated by dopamine(69) and DA dysregulation has been shown to exacerbate striatal imbalance(24). D1R agonists, which rescue cortico-striatal decoupling in our PFF mice, may act in part via FSI excitation (70). Whether DA deficiency affects cortico-striatal coupling by altering FSI function in early PD remains to be investigated.

Motor tasks requiring strong M1–striatum coordination, like digging, may thus serve as sensitive readouts for early PD. Task-specific fMRI studies in PD patients, such as finger-tapping paradigms(71–74), support this notion, revealing early abnormalities in functional connectivity. Disrupted M1–striatum coupling has been consistently reported in PD patients(8–11), with strong associations to symptom severity even at early stages(73, 75, 76). Functional decoupling is increasingly recognized as a biomarker for early neurodegenerative disease detection, including PD(77, 78) and Alzheimer’s disease(79, 80). In our study, the correlation between M1–striatum temporal alignment and digging performance in both wild-type and PFF mice underscores the behavioral relevance of this coordination.

In late-stage PD rodent models, the activity of dMSNs typically declines while iMSN activity increases, disrupting the balance between the direct and indirect pathways and contributing to motor dysfunction(45, 81). We found that even at early stages, dMSNs exhibited impaired activity during digging, whereas iMSNs were relatively preserved. This finding aligns with previous work showing selective dMSN–M1 disconnection in a dopamine-depleted PD mouse model, even in asymptomatic mice(62), indicating early vulnerability of dMSNs in PD progression. Recent studies demonstrate that gene therapy targeting dMSNs can reverse PD-like symptoms in animal models(82). Consistently, we demonstrated that administration of a D1R agonist but not D2R agonist, rescued both digging-associated decoupling and behavioral deficits. These results highlight dMSNs as promising targets for early PD intervention.

## Acknowledgments

We thank Dr. Cong Liu and his students for their help in preparing the PFF. We also thank the staff members of the animal facility at the National Facility for Protein Science in Shanghai (NFPS), Shanghai Advanced Research Institute, Chinese Academy of Sciences, China for excellent support. This work is supported by NSFC (Grant No. 32271004, 32070963), Shanghai Municipal Science and Technology Major Project (Grant No. 2019SHZDZX02), Shanghai Key Laboratory of Aging Studies (19DZ2260400) to K.-W.H.

## Author contributions

Conceptualization, K.-W.H.; Primary Investigation, X.-R.Y.; Assistant Investigation, Y.L., W.-T.Z., M.-Q.H.; Data Analysis, X.-R.Y., Y.L., K.-W.H.; Writing, X.-R.Y., K.-W.H.; Funding Acquisition, K.-W.H; Supervision, K.-W.H.

## Ethics approval

The study was approved by the Institutional Animal Care and Use Committees at the Interdisciplinary Research Center on Biology and Chemistry, Chinese Academy of Science.

## Code availability

All customized code utilized in this study is available at https://github.com/Kaiwen-Helab/Yao_etal_2024.

## Data availability

The data supporting the findings in the present study are available from the corresponding author on reasonable request.

## METHODS

### Animals

All procedures were approved by the Institutional Animal Care and Use Committees at the Interdisciplinary Research Center on Biology and Chemistry (IRCBC), Chinese Academy of Science. C57BL/6J mice were purchased from Shanghai Lingchang Biotechnology Co., Ltd.. Transgenic mice (D1R-Cre: MMRRC_034258-UCD; D2R-Cre: MMRRC_034258-UCD) were bred in-house. Mice were group housed with a 12 h:12 h light:dark cycle and ad libitum access to food and water until experiments. Male mice of 2-8 months of age were used with exception stated specifically. All experiments were carried out at ZT (Zeitgeber time) 15∼19.

### Preparation and characterization of wild-type mouse α-Syn PFF

The procedures followed the same protocol as described previously(30). In brief, pET22 vector plasmid containing α-syn gene was transfected into BL21 (DE3) cells. α-syn expression was induced by IPTG (isopropyl-1-thio-D-galactopyranoside). Cells were then harvested and sonicated followed by boiling, streptomycin (20 mg/mL) treatment, pH adjustment, and dialysis overnight in turn. The α-syn monomer was then incubated in solution containing 100 μM in 50 mM Tris, pH 7.5, 150 mM KCl, and 0.05% NaN3 buffer at 37 °C with constant agitation (900 rpm) in ThermoMixer (Eppendorf) for one week to form fibril structure. The fibril was quantified by subtracting the residual soluble α-syn from the total amount of α-syn monomer and the fibril pellet was suspended with PBS to 2 μg/μL. The obtained fibril was sonicated (20% power 15 times (1 s on, 1 s off) on ice, JY92-IIN sonicator) and the structure was confirmed by TEM (FEI Company, USA).

### Stereotaxic injection

Craniotomy was performed as described previously(30, 63). Briefly, mice were anesthetized with isoflurane vapor (1∼2% in air) and head-fixed in a stereotaxic frame (RWD Life Science, China) with heating pad (∼37 °C) underneath to maintain body temperature. The scalp was removed, and virus/fibril/PBS was delivered at a rate of 80 nl/min to either M1 (bregma/midline/depth: 1.77/1.5/-0.67 mm) or dorsal lateral striatum (dlSTR. For GCaMP7s: −0.11/2.5/2.3 mm; for PFF/PBS, DA sensor, 0.2/2.0/2.6 mm) by using a glass pipette connected to a micro syringe pump (Stoelting, USA). For photometry calcium imaging, 150 nl of AAV-hSyn-NES-jGRECO1a was injected into M1 layer V; 350 nl of AAV-hSyn-GCaMP7s (or 400nL AAV-hSyn-DIO-GCaMP7s for Cre mice) was injected into dlSTR. To measure the striatal extra-cellular dopamine level, 400 nl of AAV-hSyn-GRabeen-DA4.4 was injected into dlSTR. All viruses were diluted to a titer of 10^12^ vg./ml. For establishing the α-Syn PFF model, PFF or PBS (0.1 μl/g body weight, 2.5 μl at most for each hemisphere) was injected at a speed of 0.5 μl/min for the first 0.2 μl and 0.2 μl/min for the rest.

### Fiber photometry surgery and imaging

Optical fibers (NA: 0.37; diameter: 200μm; Inper, China) were implanted using the same M1 and dlSTR coordinates except 0.05 mm less in depth right after the virus injection. Fiber was secured to the skull using dental cement (C&M Super Bond, Japan). Mice were then housed individually and allowed to recover for at least 2 weeks before imaging.

Photometry recording during behavioral paradigms was conducted during the dark cycle utilizing a commercial multichannel fiber photometry system (Inper, China). Excitation light at wavelengths of 470 nm, 561 nm, and 410 nm was used to excite GCaMP7s/DA-4.4, RGECO1a, and Ca^2+^-independent internal channel control, respectively. Fluorescence signal was collected at 40 Hz using InperStudio (MultiColorEVAL15) software, which automatically synchronized the calcium signal with video footage from a camera positioned above the arena. Throughout the imaging, the patch cord attached to the optical fiber was suspended by a helium-filled balloon to minimize its weight impact on the mouse behavior. All behavioral trials lasted for less than 30 min.

### Immunofluorescence (IF) staining

IF was performed as previously describe(30). Briefly, mice were anesthetized with isoflurane vapor and perfused with PBS followed by 4% paraformaldehyde (PFA). Subsequently, whole brain was removed and fixed overnight in PFA at 4°C. The fixed brains were then saturated in PBS containing 20% sucrose for overnight to ensure the mice brains sinking bottom, and 30% sucrose for 2 d. Afterward, brains were embedded in tissue freezing medium (OCT, Leica) and rapidly frozen and stored at −80°C. Next, 20 μm coronal slices were sectioned using the cryostat (Leica, CM3050s-1-1-1). These sections were incubated in blocking solution (PBS containing 5% normal goat serum (Sigma) and 0.1% Triton X-100) for 2 hours at room temperature, followed by overnight incubation at 4°C with the primary antibodies diluted in the blocking solution. After incubation, the primary antibody solution was washed out three times with PBST (0.1% Tween-20 in PBS) and once with PBS, followed by incubation with the fluorophore-conjugated secondary antibodies for 2 hours at room temperature. DAPI (diluted at 1:500) was then used to stain the nuclei for 10 minutes at room temperature.

To validate the expression and location of the calcium and dopamine sensor, PFA-fixed brains were sectioned into 60 μm coronal slices using a vibratome (Leica VT 1000S). Slices with optical fiber track were selected for subsequent fluorescence imaging. Slices were further stained for either D1R or D2R to identify specific cell type.

### Confocal Imaging

Fluorescence images were captured using a spinning disk confocal microscope (Andor, UK). Different lasers with wavelengths of 630 nm, 561 nm, 488 nm, and 405 nm were sequentially utilized to excite the Fluor 647, Fluor 568 or RGECO1a, Fluor 488 or GCaMP7s or DA4.4, and DAPI respectively. Imaging was performed using a 20× air (NA = 0.75), or 40× water (NA = 1.25) objective, with consistent settings maintained for each set of experiments. The image resolution is 0.6 μm:0.6 μm/pixel (x:y) for 20× and 0.15 μm:0.15 μm/pixel (x:y) for 40× objective. Z-stack images were acquired for all brain regions and then projected in the z-direction to obtain the maximal intensity for subsequent analysis in ImageJ. Step size (μm) and thickness (μm) of each brain region were as followed: SN: 0.5, 1; Striatum:

1, 5 (for characterization of the PFF-induced pathology); Others: 2, 10. The whole view of STR was analyzed, while the SN region was manually selected based on the mouse brain atlas. The average intensities of phosphorylated α-synuclein (P-α-syn) and dopamine transporter (DAT) were measured using the same background subtraction and contrast transformation applied to each dataset, and then normalized to the control (PBS group). DAT-positive dopamine neurons in the Substantia Nigra pars compacta (SNpc) were automatically selected and counted using the Spot module of Imaris (Oxford Instruments), with settings consistent with previous descriptions(42).

### Behavioral task or test

#### a. Digging test

Mice were food deprived for 24h before behavioral digging test. During the deprivation, two food pellets (25mg, sucrose and milk powder in a 1:1 ratio. Xietong Bio-engineering Co., Ltd., China) were placed in their home cages to introduce novel food. The test environment was consisted of a lidless, semitransparent box (FENGSHI group, China), identical in size and texture to the mouse’s home cage (l:w:h: 29 cm×18 cm×13 cm), serving as the arena for the digging test. The bottom of the arena was covered with approximately 3cm-thick clean bedding, and twelve food pellets were evenly buried in the bedding. Additionally, a solitary food pellet was strategically positioned at the center of the arena’s surface on top of the bedding, easily accessible for the mice upon entering the arena, thereby enticing them to dig for buried pellets. Mice’s performances in the arena were recorded using two cameras—one providing a top view and the other a side view—for subsequent post-hoc analysis. After completing the test, the bedding was discarded, and the arena was cleaned with 75% ethanol.

#### b. Sound-cued licking task

Mice were habituated and trained for three days prior the final experiment. Day0. Completely water deprivation Day1. Mice were instructed to lick water from the water outlet connected to a sensor. Once the mouse licked the outlet, a single dose of 5 μl water was pumped out from the outlet. Each mouse repeated this process for 100 times.

Day2. A sound cue (3000 Hz, 200 ms) with random interval (10-20 s) was introduced, and mice had to associate it with water. 5 μl water was pumped out if mice licked the outlet within 3 s after the cue; otherwise, they received nothing. A 5 s interval allowed mice to consume the water before moving to the next random time interval. Each mouse was allowed to complete 80 water trials.

Day3. The random time interval from Day 2 was replaced with a 5 s non-licking interval. Mice had to be refrained from licking the outlet for 5s to trigger the sound cue. If they licked during this interval, they had to wait another 5s for the sound cue. The time window for mice to respond to the cue and to consume water remained the same as Day 2. Each mouse was allowed sufficient time to complete 120 watered trials.

Day4. Fiber photometry recording was conducted while mice performed the task, following the paradigm of Day3. The timestamp of licks within each watered trial (see Figure S5E) was recorded by a sensor controlled by customized Arduino code. The initiation of licking was defined as the first lick detected by the sensor after the sound cue.

#### c. Rotarod

The procedures followed the protocol described in a previous report(30). Briefly, mice underwent three days of training, consisting of two adaptive trials at a low rolling speed of 5 rpm/min, followed by a third trial with accelerating speed ranging from 0 to 40 rpm/min. After a rest day on the fourth day, mice were tested twice on the rotarod on the fifth day. Each testing session lasted for a total of 5 minutes. During the first 1.5 minutes, the speed of the rotarod accelerated from 0 to 40 rpm/min, which was maintained for the remaining time. Mice were tested twice, and the maximum duration they stayed on the rotarod was recorded for final analysis. Between each trial, the rotarod was thoroughly cleaned with 75% alcohol.

#### d. Open field test

After a period of habituation, mice were introduced into the center of the open field chamber (40 × 40 cm) and given 10 mins to freely explore its surroundings. After the test, the arena was cleaned with 75% ethanol. Their activities were observed and recorded using a camera positioned on the ceiling, and the data were subsequently analyzed using EthoVision XT 11.5 software (Noldus, Netherlands).

#### e. Free exploration of a new cage

The procedures and settings for the open field test were replicated with the exception of utilizing a lidless and empty cage resembling to the mouse’s home cage (l:w:h: 29 cm×18 cm×13 cm) as the arena.

#### f. Free moving

The procedures and settings were exactly the same with free exploration but the arena covered with clean bedding.

#### g. Treadmill running

Mouse was placed on the treadmill (Zhenghua Biologic apparatus, China) 10 mins before testing for habitation. During the test, the rolling velocity of conveyor was set at constant 10 m/min and the testing session lasted for 10 mins. To prevent interaction between mice, each mouse was isolated by baffles while running on the treadmill. After the test, conveyor and baffles were cleaned with 75% ethanol. Both male and female mice were included in this experiment but they were separated by gender during the test. Mice that failed to keep up with rolling conveyor during the test was excluded from the analysis.

### Pharmacology

The experimental design followed a previous study that investigated the effects of dopaminergic drugs on striatal MSNs using calcium imaging in parkinsonian mice(26). Drugs (all compounds from Merck) including D1R agonist (SKF81297; 1 mg/kg), D2R agonist (quinpirole; 0.5 mg/kg), L-DOPA (1 mg/kg) together with benserazide (15 mg/kg), were dissolved in 0.9% saline and filtered through a PES membrane filter unit (0.22 μm, Millex). These solutions were stored at 4℃ and used within a week. Drugs were administered via intraperitoneal (i.p.) injection, with a volume determined according to body weight (0.1 ml/10 g). Both behavioral and photometry experiments commenced 15 minutes after i.p. except quinpirole (60 minutes post i.p.).

### Optogenetics

400 ul of AAV-hSyn-ChR2-GFP or AAV-hSyn-GFP at a titer of 10^12^ vg./ml were bilaterally injected into M1 and STR using the same coordinates as those used for photometry recording. Optical fibers were then implanted 200 μm above each injection site. Following a four-week recovery period, mice underwent digging test (see Methods: Digging Test), during which 470nm blue light pulses (frequency: 5 Hz, duration: 5 ms, intensity: 5 mW) were delivered bilaterally to M1 and striatum using a commercial optogenetic instrument (Aurora-300, NEWDOON, China) following pre-defined patterns. Four stimulation patterns were tested, each lasted for 6 min: 1) synchronized stimulation: identical light pulses were delivered simultaneously to M1 and striatum; 2) desynchronized stimulation: 50 ms delay was introduced between M1 and striatal stimulation; 3) M1-only stimulation; 4) STR-only stimulation. The order of synchronized and desynchronized stimulations was counterbalanced across animals to control for order effects.

### Quantification and statistics

#### Behavioral analysis

Digging behavior. Trials were manually picked from the video by the investigator. The initiation of digging behavior was determined when the bedding beneath the paws of the mice were visibly displaced. The termination of a digging trial was defined as either no bedding displacement for 1 s or the mouse started engaging in any other behaviors such as rearing, grooming, or eating pellets. The average duration of all digging events within the first ten minutes after mice acquired the center pellet on top of the bedding was considered as the digging duration. Digging frequency was defined as the number of digging per minute during this period. The success rate was calculated as the percentage of pellet-retrieving digging events out of the total.

Licking behavior. The whole experiment was controlled and recorded by customized Arduino code. Each lick was automatically captured and timestamped. Data was analyzed by customized R code. A series of successive licks with intervals less than 1 s was considered as a bout. The duration of a bout was defined as the bout length.

Contralateral turning behavior. Trials were manually picked from the video by the investigator. Only turning behaviors exceeding 90 degrees and not overlapping with digging were included in the analysis.

#### Qualitative analysis of photometry signal

##### a. Photometry signal processing

The photometry data collected was pre-processed for bleaching and motion correction using InperDataProcess v0.7.2 software. The onset of digging and contralateral turning behavioral events was manually marked based on the synchronized videos. For digging behavior, only trials having ≥ 10 s of non-digging baseline and ≥ 3 s of digging were selected for final analysis. For turning behavior, only trials with ≥ 4 s inter-trial interval and ≥ 90 degree turning were chosen for final analysis.

All fiber signal was presented as ΔF/ F_0_ using the equations:

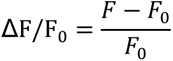

F represents the fluorescence value of each time-point. F_0_ is the average of F of baseline.

For digging and licking behaviors, the time window for analysis was −5 to 5 s relative to behavior initiation, during which the −4 to −2 s was used to calculate the F_0_. For contralateral turning, the time window was −1 to 1 second, and the baseline was -1 to 0 s.

##### b. Correlation coefficient (static and sliding window) & Phase lag

The static (global) correlation coefficient was calculated by customized python code: the data (ΔF/ F_0_) was bandpass filtered (0.001∼3Hz) at first, then trimmed 1 sec both at the beginning and ending (for licking and digging behavior only). The Pearson’s coefficient was calculated by corrcoef function of numpy.

To assess the dynamic coordination between the two calcium signals over time, time-resolved correlation coefficients was computed using a sliding window approach. A window of 4 sec was advanced in 0.5-sec increments across the whole analyzed epoch. Within each window, Pearson’s correlation coefficient (*r*) was calculated to quantify the linear relationship between the two traces. Edge effects at the end of the recording period were addressed by excluding incomplete windows from final analysis.

The phase lag was then identified as the lag value at which the absolute cross-correlation function reached its maximum, indicating the optimal temporal shift for signal alignment (see Fig. 4B), and the corresponding time lag was determined with signal.correlation_lags function of SciPy.

##### c. Plateau & peak amplitude

The plateau amplitude was defined as the mean of the normalized (ΔF/F_0_) signal within the plateau stage (1∼2 s relative to initiation of digging), or peak (0∼0.5 s relative to the initiation of licking).

##### d. Sum dFF activity

The data of each mouse collected during 10 minutes of treadmill running or free-moving was separated into bins of equal length, typically 8 seconds each. Each bin of data was normalized to ΔF/F_0_, where F_0_ represented the 20^th^ percentile of the data within that bin. In the meanwhile, we calculated the Pearson’s correlation coefficient between the derived ΔF/F_0_ and the raw F. Data with lower than 0.8 correlation coefficient was excluded from final analysis. The sum dFF activity was then determined by summing the ΔF/F_0_ values across all bins.

### Statistical analysis

Sample sizes were identified in all figures. Statistical analysis was conducted using Prism V9.0 software (GraphPad Software, Inc.). For unpaired data, the non-parametric Mann-Whitney (M-W) test was employed. Wilcoxon matched-pairs test was used for paired comparison. One-way balanced ANOVA followed by Šídák’s or Dunnett’s multiple comparison test was used for comparisons among multiple groups. Cumulative distributions were compared using the Kolmogorov-Smirnov (KS) test. The significant level was set at P < 0.05, denoted as *P < 0.05, **P < 0.01, and ***P < 0.001. Additionally, when the P-value fell between 0.05 and 0.1, it was labeled accordingly. Error bars in all figures represent the standard error of the mean (s.e.m).

## Table of materials

| Antibodies | Host species | Company | Identifier | Dilution |
| --- | --- | --- | --- | --- |
| Phospho-α-syn (S129) | rabbit | Abcam | ab51253 | 1:250 |
| DAT | rat | Abcam | ab5990 | 1:500 |
| Alexa Fluor 568 | goat (anti-rabbit) | Abcam | ab175471 | 1:1000 |
| Alexa Fluor 647 | goat (anti-rat) | Abcam | ab150159 | 1:150 |
| ChAT | rabbit | Proteintech | 20747-1-AP | 1:400 |
| DRD1 | rabbit | Proteintech | 17934-1-AP | 1:200 |
| DRD2 | rabbit | Proteintech | 55084-1-AP | 1:200 |
| Alexa Fluor 647 | goat (anti-rabbit) | Thermo Fisher | A21245 | 1:1000 |
| Chemicals | Product/chemical name | Company | CAS number | Note |
| SKF-81297 | 6-Chloro-PB hydrobromide | Sigma-Aldrich | 71636-61-8 | D1R agonist |
| Quinpirole | Quinpirole Hydrochloride | Sigma-Aldrich | 85798-08-9 | D2R agonist |
| L-dopa | Levodopa | Sigma-Aldrich | 59-92-7 | Precursor of dopamine |
| Benserazid e | Benserazide hydrochloride | Sigma-Aldrich | 14919-77-8 | Combined with L-dopa |

## Supplementary figures and legends

**Figure S1.**
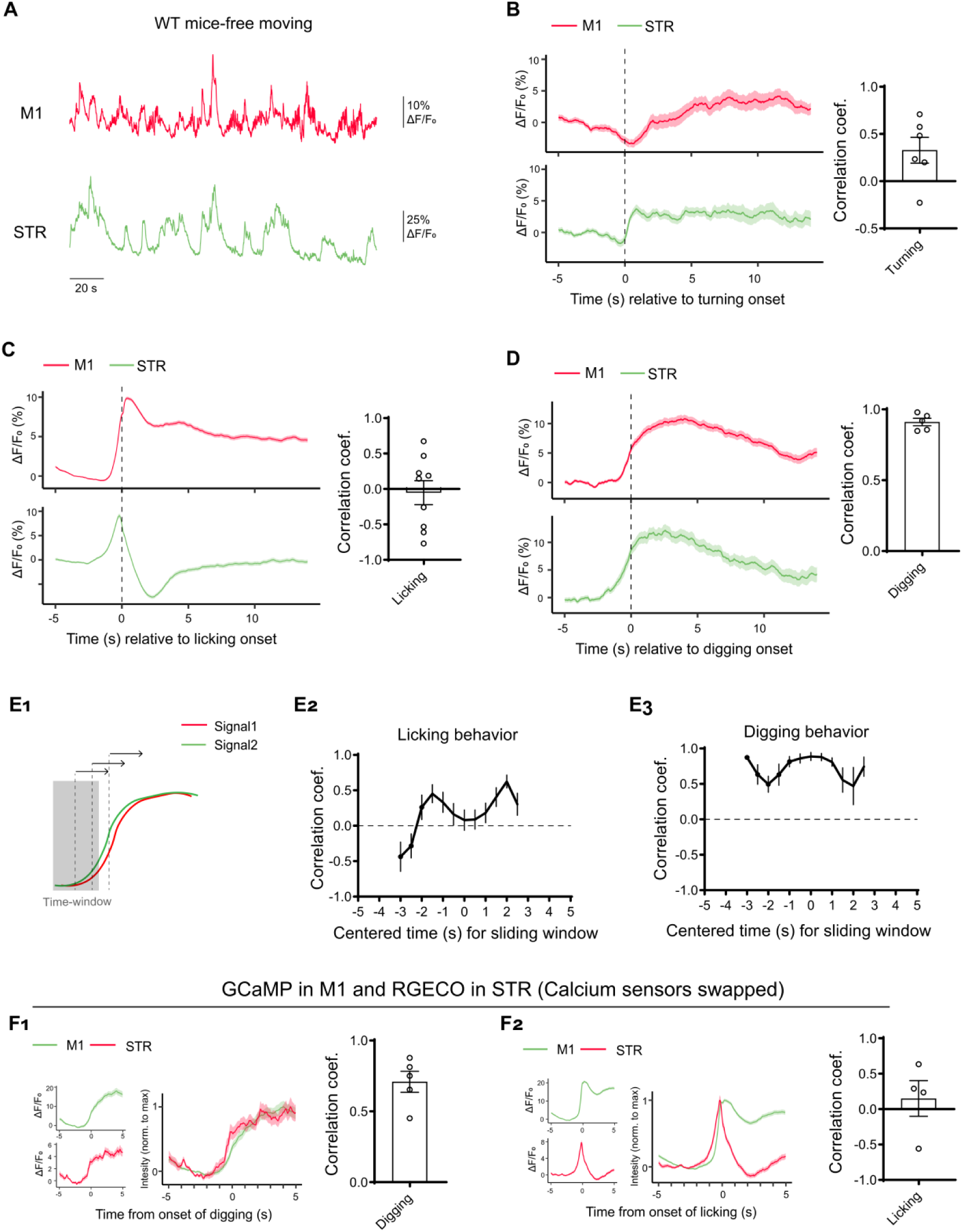
cortico-striatal coupling was behavior-specific and independent of calcium sensors. **A.** Example temporally aligned Ca^2+^ activities recorded from M1 (red) and STR (green) from a wild-type mouse free moving in a lidless cage. **B-D**. Prolonged time-window recordings of calcium activity in M1 and striatum (STR). (Left panel) Temporally aligned calcium traces (ΔF/F₀) for M1 and STR across contralateral turning (B), licking (C), and digging (D) behaviors in wild-type mice. (Right panel) Cross-correlation analysis of M1-STR calcium signals during each behavior. **E.** Time-resolved cross-correlation analysis of M1 and striatal Ca²⁺ activities using a 4-second sliding window. (E_1_): Schematic depiction of analysis. (E_2_): Analysis for licking. (E_3_): Analysis for digging behavior. **F**. Normalized ΔF/F₀ traces of M1 (green) and STR (red) temporally aligned to digging-associated (F_1_) or licking-associated (F_2_) behaviors, along with the corresponding correlation coefficient, from wild-type mice injected with region-reversed calcium indicators.

**Figure S2.**
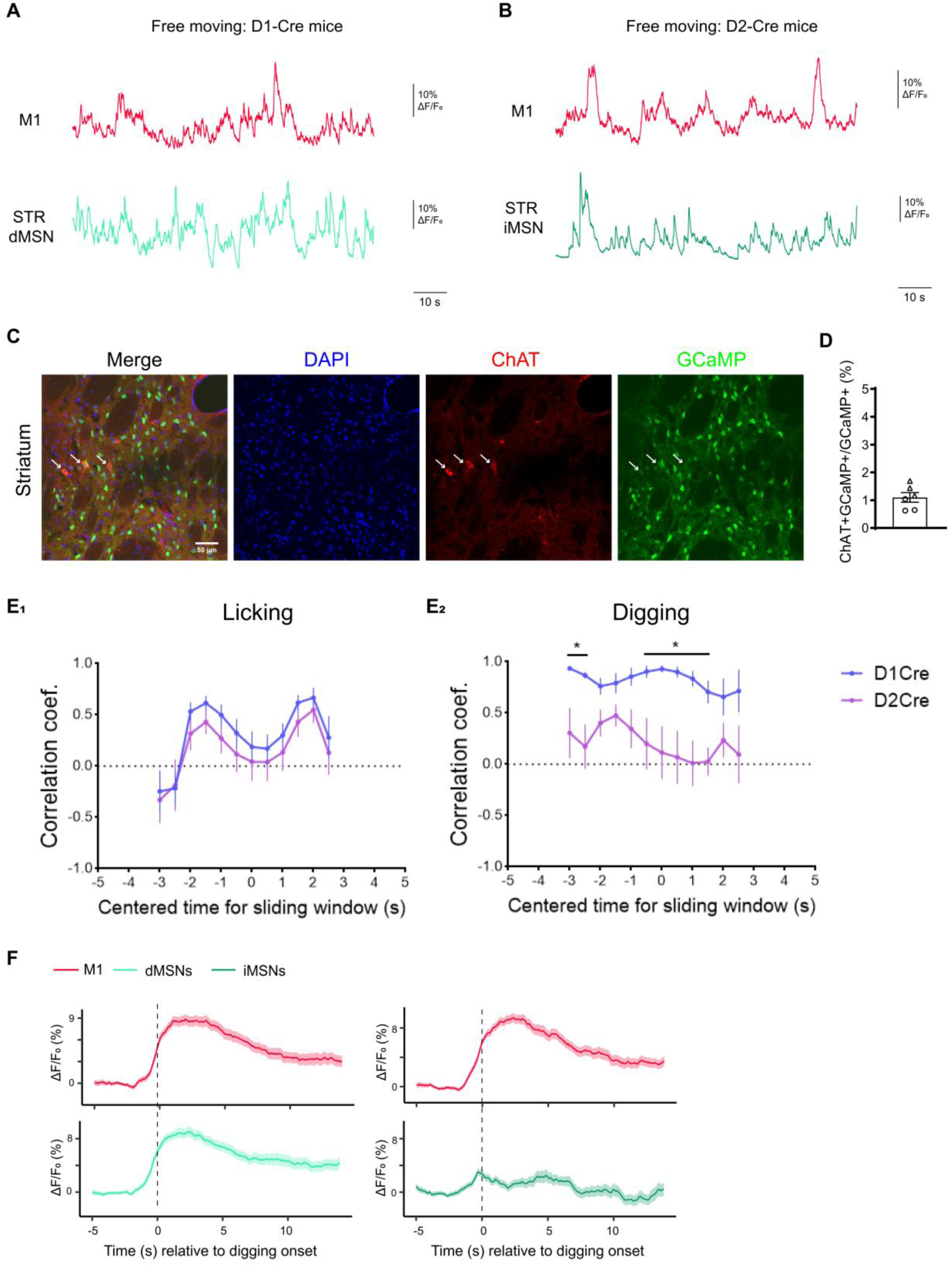
Validation of cell type-specific resolution in the methodology. Related to Fig2. **A**. Example temporally aligned Ca^2+^ activities recorded from M1 (red) and STR dMSNs (aqua) from a D1R-Cre mouse during free moving. **B**. Example temporally aligned Ca^2+^ activities recorded from M1 (red) and STR iMSNs (dark green) from a D2R-Cre mouse during free moving. **C-D.** Co-immunostaining of ChAT and GCaMP7 in coronal brain slices harvested from D2-cre mice expressing DIO-GCaMP in the striatum. (C) Representative images. Scale bar: 50 μm. (D) Quantification of the percentage of ChAT+GCaMP+ cells relative to all GCaMP+ cells. **E.** Comparison of correlation coefficient of M1 and striatal Ca^2+^ activities within sliding window (length: 4s) from D1-Cre mice (blue) and D2-Cre mice (purple) for licking (E_1_) and digging behavior (E_2_). Two-way ANOVA with Sidak’s multiple comparisons test. *: *P* < 0.05. **F.** Prolonged time-window recordings of digging associated calcium activity (ΔF/F₀) of M1 and striatal dMSNs (light green) or iMSNs (dark green) from D1R-Cre or D2R-Cre mice.

**Figure S3.**
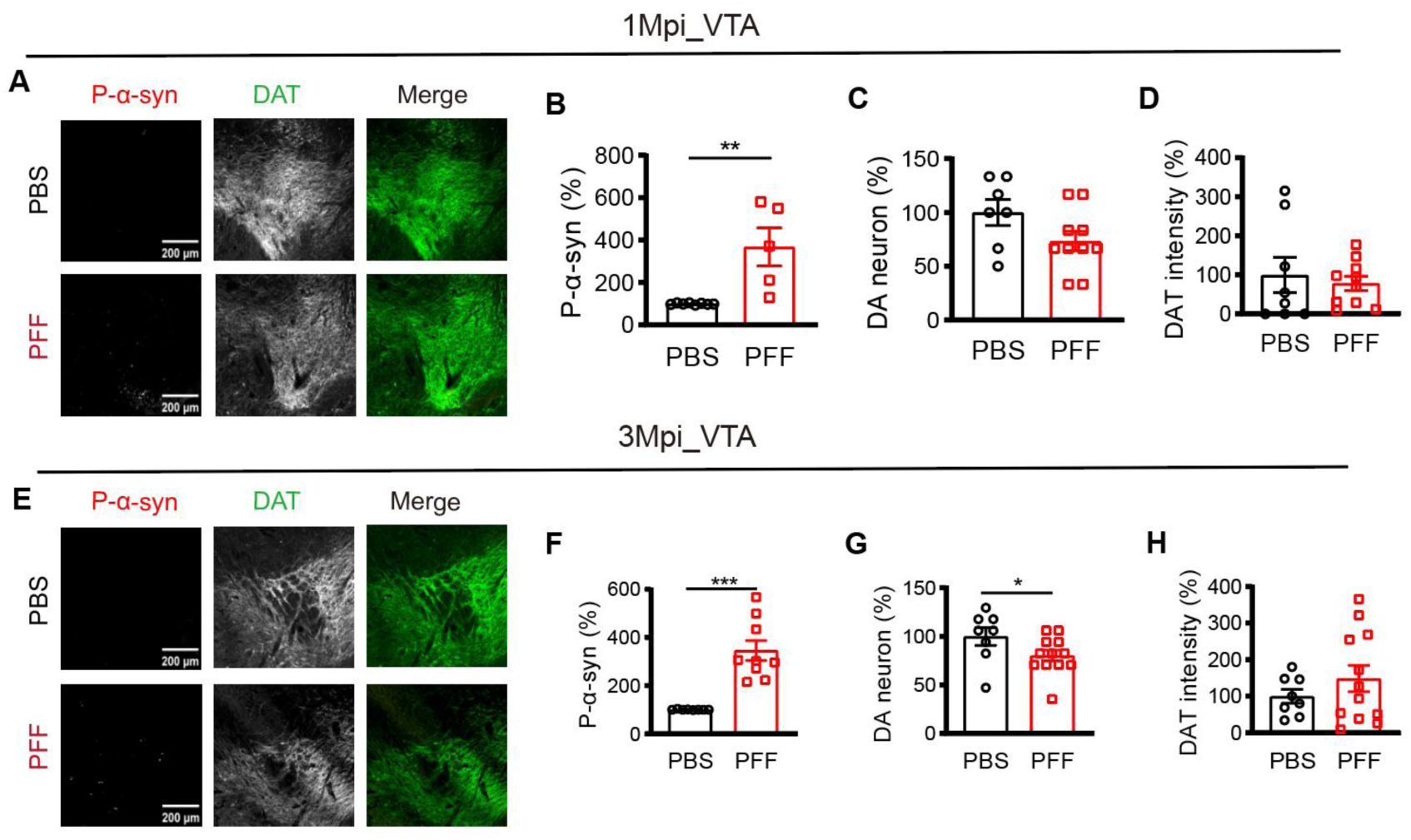
Pathologies in VTA region of PD-like mice. Related to Fig3. **A-D.** Immunofluorescence analyses of pathological hallmarks in VTA regions in PFF- and PBS-injected mice at 1 Mpi. A, Representative immunofluorescence images. Scale bars: 200 μm. (B-D) Comparisons of the p-α-Syn intensity (B, *P* = 0.0016), DA neuron number (C) and DAT intensity (D) in VTA. All data were normalized to PBS control (PBS: 8 views from 2 mice; PFF: 10 views from 3 mice). **E-H.** Immunofluorescence analyses of pathological hallmarks in VTA regions in PFF- and PBS-injected mice at 3 Mpi. All data normalized to PBS control. E, Representative immunofluorescence images. Scale bars: 200 μm. (F-H) Comparison of the p-α-Syn intensity (F, *P* < 0.0001), DA neuron number (G, *P* = 0.0368) and DAT intensity (H) in VTA (PBS: 8 views from 2 mice; PFF: 12 views from 2 mice).

**Figure S4.**
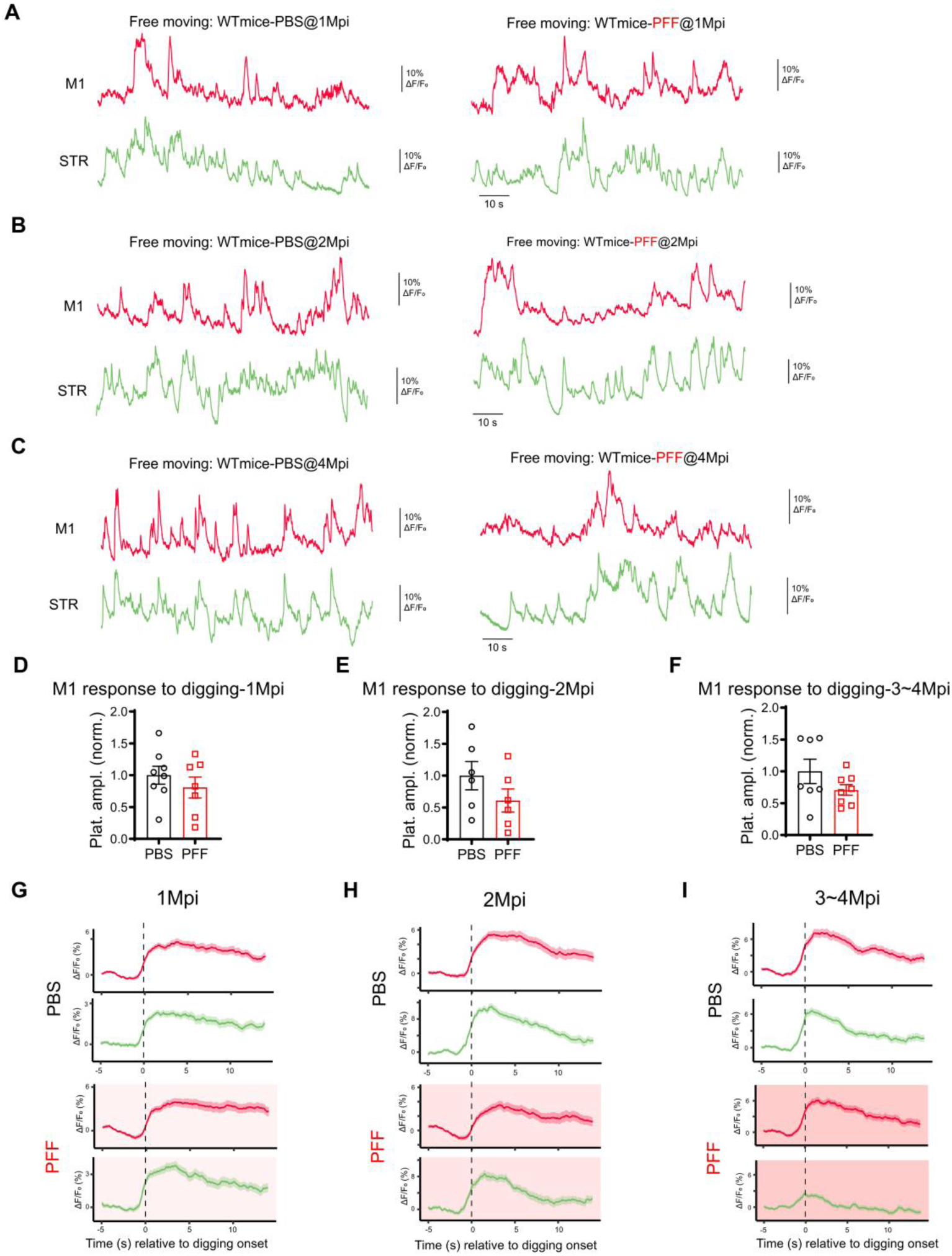
Cortico-striatal decoupling in PFF mice was behavior-specific. Related to Fig4. **A-C**. Example temporally aligned Ca^2+^ activities recorded from M1 (red) and STR (green) from a PBS-(left) or PFF-injected mouse (right) during free moving at 1 Mpi (A), 2 Mpi (B), 4 Mpi (C). **D-E**. Normalized plateau amplitude of M1 digging-associated Ca^2+^ activities from PBS- and PFF-injected mice at 1 Mpi (D), 2 Mpi (E), 3∼4 Mpi (F). Data batch-normalized to the PBS group. Mann-Whitney test. **G.** Prolonged time-window recordings of digging associated calcium activity (ΔF/F₀) of M1 and striatum from PBS or PFF mice at 1 Mpi. **H.** Prolonged time-window recordings of digging associated calcium activity (ΔF/F₀) of M1 and striatum from PBS or PFF mice at 2 Mpi. **I.** Prolonged time-window recordings of digging associated calcium activity (ΔF/F₀) of M1 and striatum from PBS or PFF mice at 3-4 Mpi.

**Figure S5.**
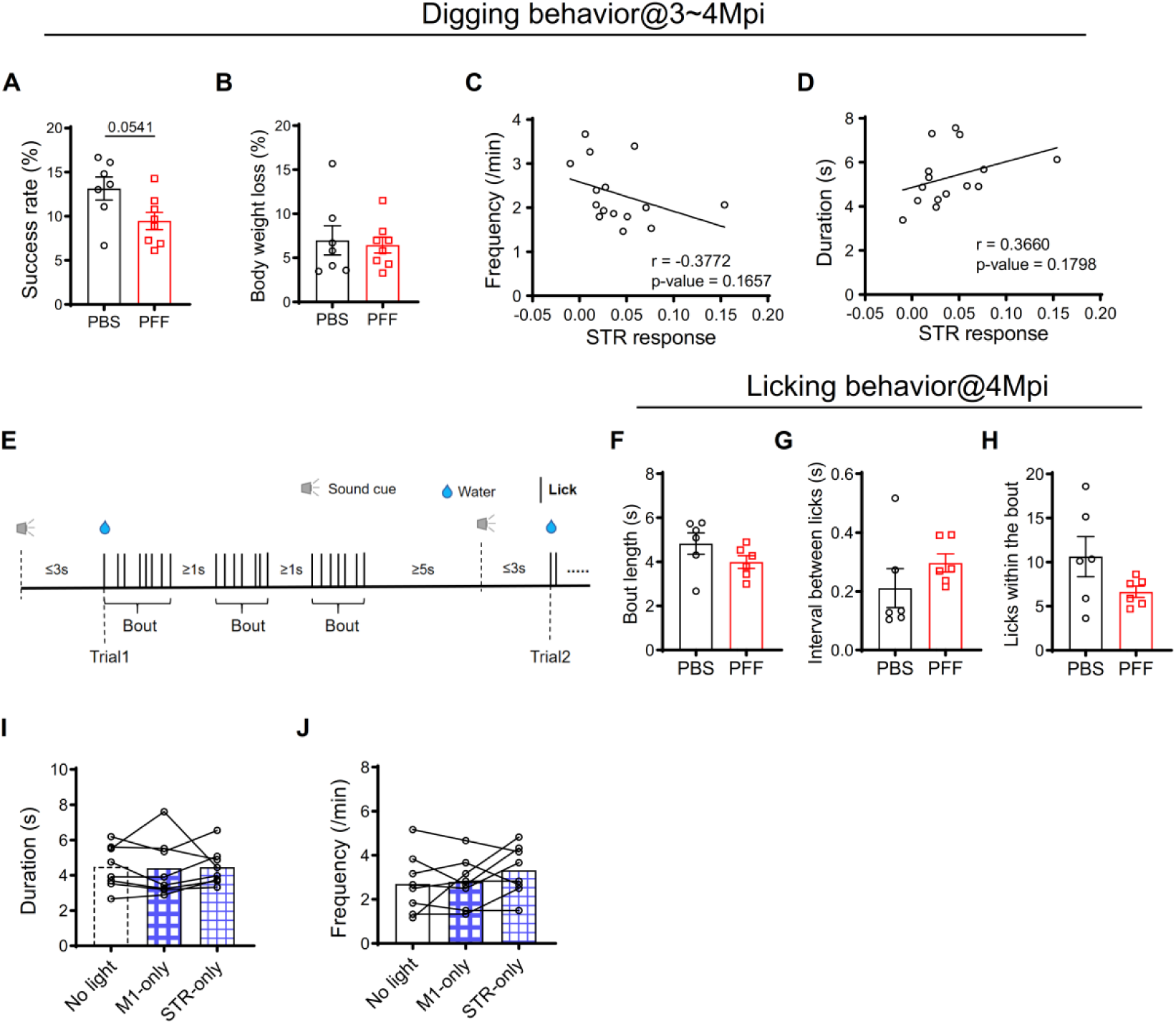
Behavioral phenotypes in PBS- or PFF-injected and ChR2-expressing wild-type mice. Related to Fig5. **A.** Success rate of digging (Number of trials retrieving pellets / Number of total trials * 100%). **B.** Body weight loss after 24h food deprivation. **C-D**. Correlation analysis between STR responses (plateau amplitude, ΔF/F₀) and digging duration (C) or frequencies (D). r, Pearson’s correlation coefficient. **E**. Paradigm of sound-cued licking test. See details in methods. **F-H**. Characterization of licking behavior. (F) Average bout duration. (G) Average lick interval within a bout. (H) Average lick counts per bout. **I.** Diagrams depicting the optogenetic stimulation protocols: three 6-minute epochs: no light stimulation, stimulation of M1 alone (M1-only) and stimulation of STR alone (STR-only). **J**. Optogenetic stimulation of M1 or STR alone did not affect digging duration (K_1_) or total digging events (K_2_). One-way ANOVA followed by Šídák’s multiple comparisons test. Mann-Whitney test.

**Figure S6.**
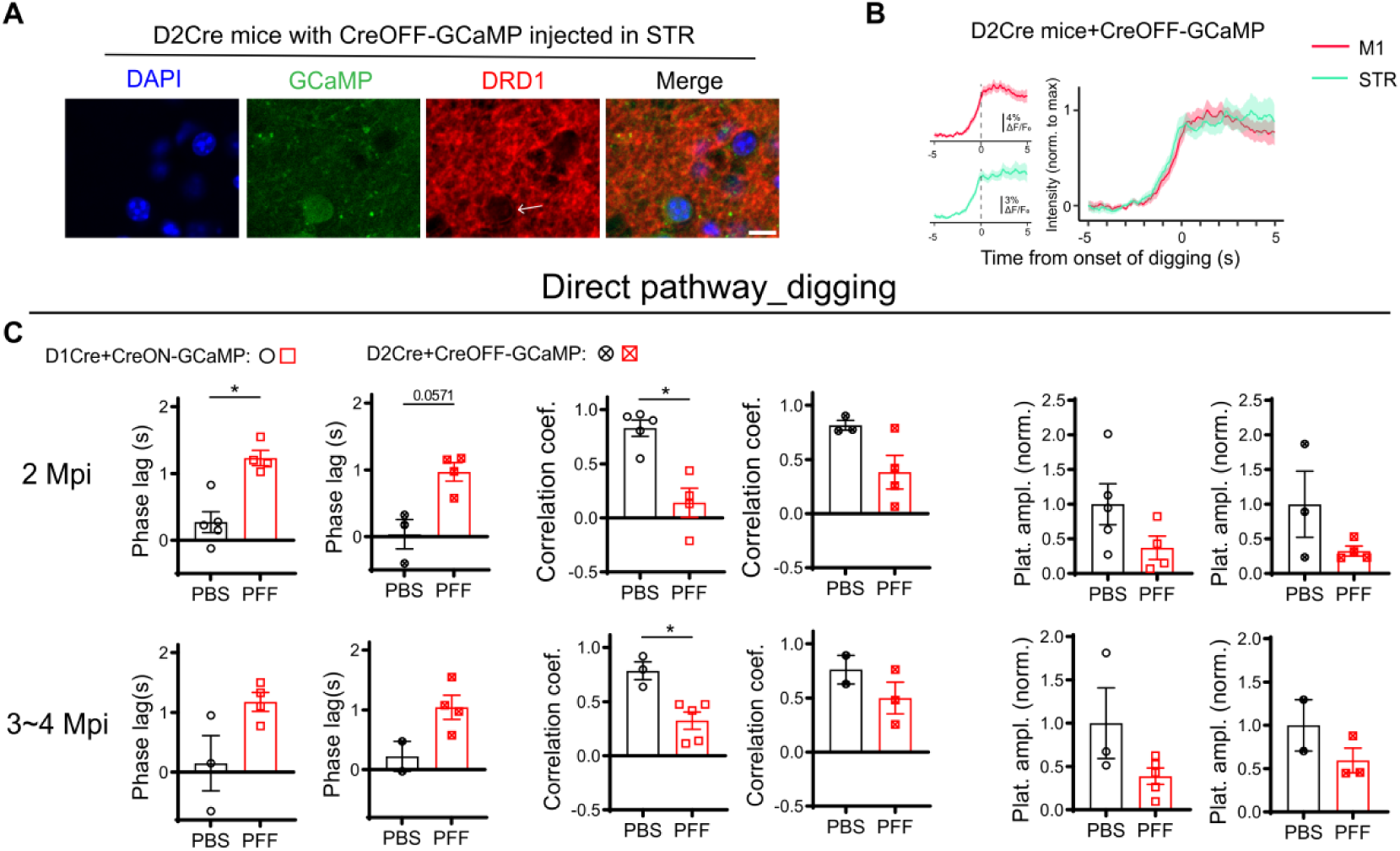
CreON and CreOFF systems equally enabled detection of the abnormal M1- dMSNs coupling associated with digging in PFF mice. Related to Fig6. **A**. Representative immunofluorescence images showing the co-localization of GCaMP and D1R immunoreactivity (arrow) from a D2R-Cre mouse expressing Cre-OFF GCaMP. Scale bar: 10 μm **B**. ΔF/F₀ traces of M1 (red) and STR dMSNs (aqua) Ca^2+^ activities temporally aligned to the initiation of digging in 3 D2R-Cre mice injected with CreOFF-GCaMP. The average ΔF/F₀ waveform of M1 (Top left panels) and STR (Bottom left panels) were presented, and the waveform for both regions were overlaid after normalizing each channel to its maximum value (panels on the right). **C.** M1-dMSNs correlation data collected from D1R-Cre mice using Cre-on GCaMP (Open symbols) and D2R-Cre mice with Cre-off GCaMP (Cross-filled symbols). Mann-Whitney test.

**Figure S7.**
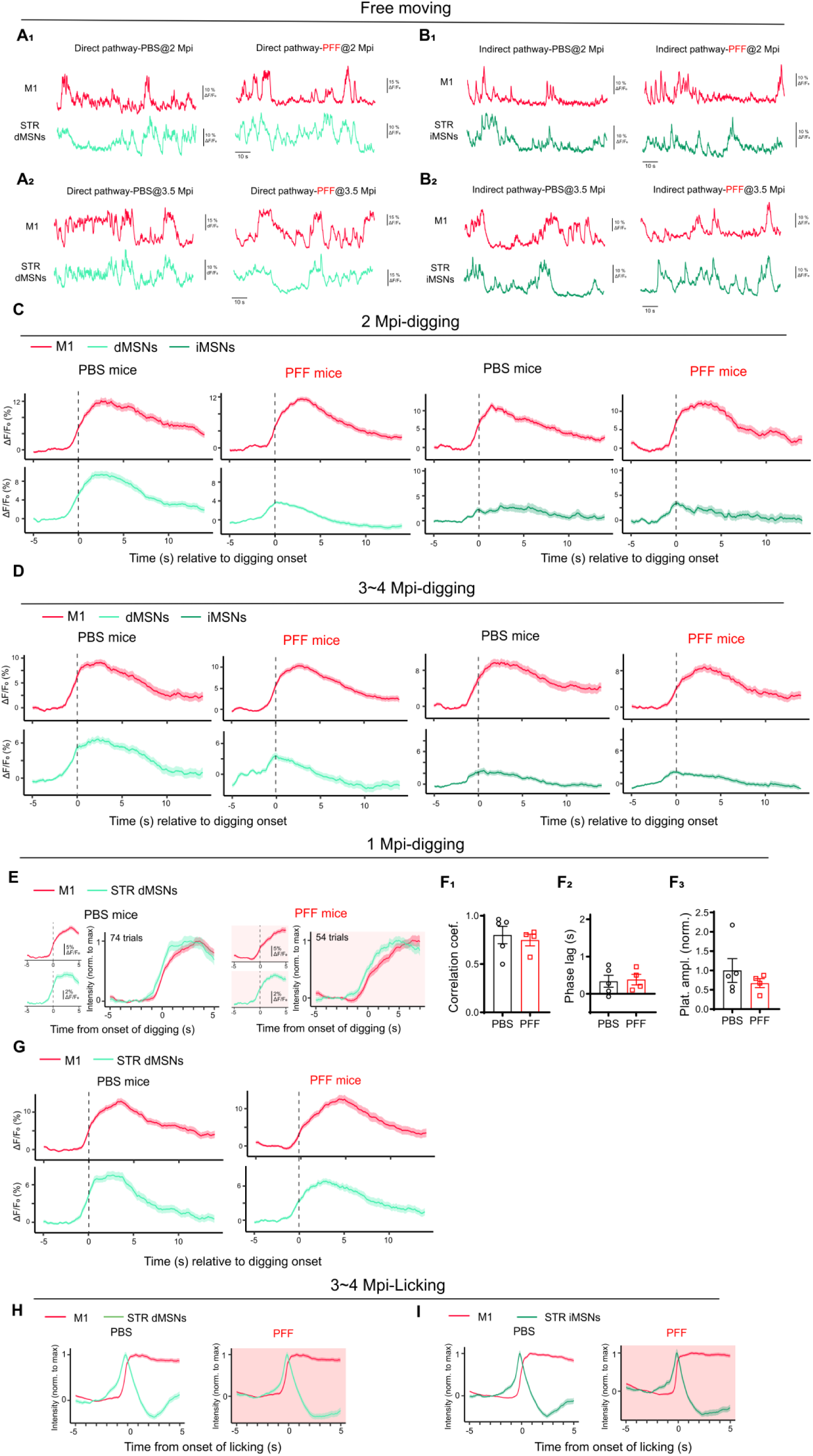
Ca^2+^ activities traces recorded from M1 and STR in behavior- and cell type-specific manner. Related to Fig6. **A.** Example temporally aligned Ca^2+^ activities traces recorded from M1 (red) and STR dMSNs (aqua) of PBS- (left) or PFF-injected mice (right) during free moving at 2 Mpi (A_1_) and 3.5 Mpi (A_2_). **B.** Example temporally aligned Ca^2+^ activities traces recorded from M1 (red) and STR iMSNs (dark green) of PBS- (left) or PFF-injected mice (right) during free moving at 2 Mpi (B_1_) and 3.5 Mpi (B_2_). **C.** Prolonged time-window recordings of digging associated calcium activity (ΔF/F₀) of M1 and striatal dMSNs (light green) or iMSNs (dark green) from PBS or PFF mice at 2 Mpi. **D.** Prolonged time-window recordings of digging associated calcium activity (ΔF/F₀) of M1 and striatal dMSNs (light green) or iMSNs (dark green) from PBS or PFF mice at 3∼4 Mpi. **E-G**. Dual-site fiber photometry imaging of M1 and striatal dMSNs in PFF- or PBS-injected mice at 1 Mpi (PBS: 74 trials from 5 mice; PFF: 54 trials from 4 mice) during digging. (E) Normalized ΔF/F₀ traces of M1 (red) and striatal dMSNs (green) Ca^2+^ activities temporally aligned to the initiation of digging at 1 Mpi in mice injected with either PBS (panels with white background) or PFF (panels in pink background). The average ΔF/F₀ waveform of M1 (Top left panels) and STR (Bottom left panels) were presented, and the waveform for both regions were overlaid after normalizing each channel to its maximum value (panels on the right). (F) Quantitative analyses of the M1-dMSNs correlation and normalized plateau amplitudes of dMSN response at 1 Mpi. (G) Prolonged time-window recordings of digging associated calcium activity (ΔF/F₀) of M1 and striatal dMSNs (light green) from PBS or PFF mice at 1 Mpi. **H-I**. Normalized ΔF/F₀ traces of M1 (red) and STR dMSNs (H, PBS: n = 131 trials from 3 mice, PFF:133 trials from 3 mice) or iMSNs (I, 124 trials from 3 mice, PFF:162 trials from 4 mice) licking-associated Ca^2+^ activities from PBS- and PFF-injected mice at 3∼4 Mpi.

**Figure S8.**
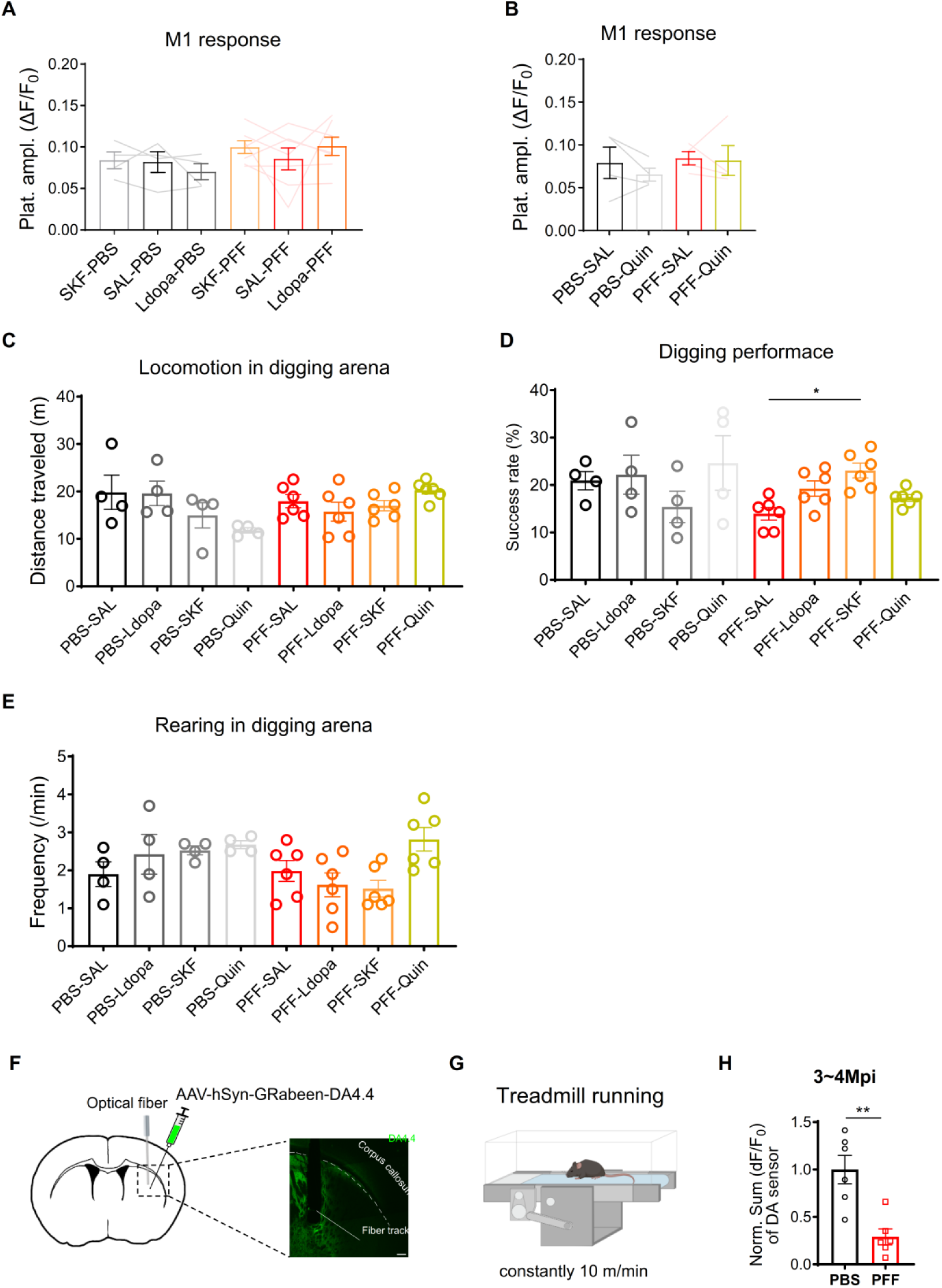
Effects of dopaminergic drugs on M1 responses and digging behavior. Related to Fig7. **A-B**. Plateau amplitudes of digging-associated M1 Ca^2+^ activities of PBS- and PFF-injected mice after administration of different drugs. **C.** Travel distances during the first 10 minutes after administration of different drugs. **D.** Effects of different drugs on digging success rates. (One-way ANOVA, F_(2.031, 10.16)_ = 6.532. SAL-PFF *vs.* SKF-PFF, *P* = 0.0478, Dunnett’s multiple comparisons test). **E.** Frequencies of rearing during the first 10 minutes after administration of different drugs. Wilcoxon matched-pairs test (B). **F.** Schematic diagram showing the experimental design for assessing striatal extracellular dopamine (DA) level. DA4.4 is a florescence-based DA sensor. Right panel, anatomical verification of virus expression and optic fiber localization. Scale bar: 200 μm. **G.** Schematic diagram for treadmill running test. **H.** Normalized sum activity of DA sensor during treadmill running from PBS- and PFF-injected mice at 3∼4 Mpi (*P* = 0.0043). Mann-Whitney test.

